# A PIWI protein-dependent DNA N6-adenine methylation pathway in *Oxytricha* protects genomic sequences from deletion

**DOI:** 10.64898/2026.01.06.698049

**Authors:** Margarita T. Angelova, Yi Feng, Danylo J. Villano, Erhan Aslan, Michael W. Lu, Pierre-Olivier Estève, Sagnik Sen, William E. Jack, Sriharsa Pradhan, Laura F. Landweber

## Abstract

The ciliate *Oxytricha* undergoes massive genome rearrangements during development to produce a functional nucleus from an encrypted zygotic genome. PIWI-interacting small RNAs protect DNA regions against deletion, but how that protective mark is established has been a mystery. Recently our lab discovered MTA1, a methyltransferase that catalyzes DNA N6-adenine (6mA) methylation. Both MTA1 and the *Oxytricha* Piwi protein, Otiwi1, are required for development, and Otiwi1 mutation eliminates 6mA signal. To examine the role of 6mA, we analyzed its genome-wide distribution across development in wild-type and MTA1 mutant backcrossed cells. We find specific and abundant enrichment on retained sequences, suggesting a protective role for this epigenetic mark. Furthermore, programmed retention of a DNA region that is normally deleted leads to accumulation of new 6mA marks on the ectopically retained DNA sequence. Together, these results suggest that piRNA-guided 6mA DNA methylation leads to protection of DNA sequences against deletion during nuclear differentiation.

**Highlights:**

1. DNA N6-methyladenine accumulates on retained DNA regions during *Oxytricha* development.
2. *mta1* mutant backcrosses have disrupted DNA methylation and a developmental delay.
3. DNA N6-adenine methylation during genome rearrangement requires the presence of Otiwi1.
4. Programmed retention of a germline-limited region leads to developmental methylation.

## Introduction

Classic epigenetic mechanisms, such as DNA methylation, histone modifications, non-coding RNAs, and nucleosome positioning, regulate epigenetic inheritance and genome-environment interactions. Adenine DNA methylation, specifically N6-methyladenine (6mA), while abundant in prokaryotes and controversial in some metazoa^1–6^, is a conserved epigenetic mark implicated in diverse cellular features of microbial eukaryotes, especially chromatin remodeling and transcription^7–15^. In green algae, basal fungi, and many ciliates, 6mA is often enriched in symmetrically dimethylated ApT motifs within nucleosome linker regions, particularly near promoters^7,9,12,13,15–21^. While established as an epigenetic mark in these representative lineages, its potential roles in development and differentiation are just beginning to emerge.

Nuclear differentiation in ciliate genera such as *Oxytricha, Tetrahymena,* and *Paramecium* offer ideal model systems to study the roles for DNA methylation in development. After cells conjugate, a transcriptionally-active somatic macronucleus (**MAC**) develops from a copy of the zygotic, germline micronucleus (**MIC**)^22^. In *Oxytricha trifallax*, this precursor-product maturation involves large-scale natural genome editing that eliminates most (>90%) of the MIC genome, including precise deletion of over 225,000 *internal eliminated sequences* (**IES**s), together with rearrangement and sometimes unscrambling of the remaining exon-sized *MAC-destined sequences* (**MDS**s)^23–27^. MDSs are typically joined at short direct repeats called ***pointers*** that define MDS/IES borders^23,24^ but are insufficient to program accurate junction formation, which requires maternally-inherited long, template RNAs^28,29^. Both retained and deleted sequences are often much smaller than the size of a nucleosome unit^23^, which led us to investigate base modifications^7,30^. The entire 3-day process of development leads to production of over 16,000 unique functional “nanochromosomes” in the MAC, most of which encode single genes and are amplified to high copy number^23,27^. We previously showed that a transgenerationally-inherited class of 27nt small RNAs (piRNAs)], associated with the PIWI-subfamily protein Otiwi1, are necessary and sufficient to mark and protect the short retained regions on precursor MIC chromosomes^31,32^. However, *Oxytricha* piRNA abundance peaks in early conjugation and declines before IES elimination and MDS rearrangement take place^31^. Thus it has been a mystery as to how *Oxytricha* piRNAs transmit their protective signal to the developing genome.

We previously identified and biochemically and genetically characterized the MTA1 DNA N6-adenine methyltransferase in *Oxytricha trifallax* and *Tetrahymena thermophila*^7^. It is part of a 4-protein complex, MTA1c, composed of two MT-A70 domain-containing proteins: MTA1 (also called AMT1 in *T. thermophila*) and MTA9 (AMT7)—plus two homeobox-domain-containing proteins that likely contribute to substrate binding^33–35^ and permitted an evolutionary switch to methylation of DNA instead of RNA^7,35^. Notably, MTA1 disruption in *Oxytricha* causes lethality during development^7^, underscoring its importance and candidate participation in genome rearrangement. Here, we demonstrate a functional role of 6mA during *Oxytricha* genome reorganization, when 6mA is naturally abundant. We find it is specifically enriched on retained DNA regions in the developing genome. Our data suggest a model in which 6mA acts as an epigenetic signal that leads to protection of genomic regions against DNA deletion during genome remodeling. Furthermore, 6mA colocalizes with and is dependent on Otiwi1, suggesting a direct interplay between the piRNA pathway and adenine DNA methylation. These findings demonstrate a developmentally critical function for 6mA, positioning it as a key epigenetic signal during RNA-guided genome editing in *Oxytricha,* and providing the missing link between piRNA marking of genomic regions for protection and their rearrangement later in development.

## Results

### Adenine DNA methylation levels peak during development

To capture the dynamics of N6-adenine DNA methylation during programmed genome rearrangement, we examined 6mA levels across post-zygotic development and nuclear differentiation in *Oxytricha trifallax*. Immunofluorescence analysis revealed a strong 6mA signal specific to the developing nucleus 24h post-mixing of mating-compatible wild-type strains JRB310^25^ and JRB510^36^, suggesting a possible role in DNA rearrangement (Figure 1A). 6mA levels continue to increase during development, peaking at 36–60h, with immunofluorescence quantification confirming this temporal pattern (Figure 1B). DNA mass spectrometry independently validated the sharp increase in the fraction of methylated deoxyadenosines, with 6mA peaking at 48h (Figures 1C and S1A).

**Figure 1.**
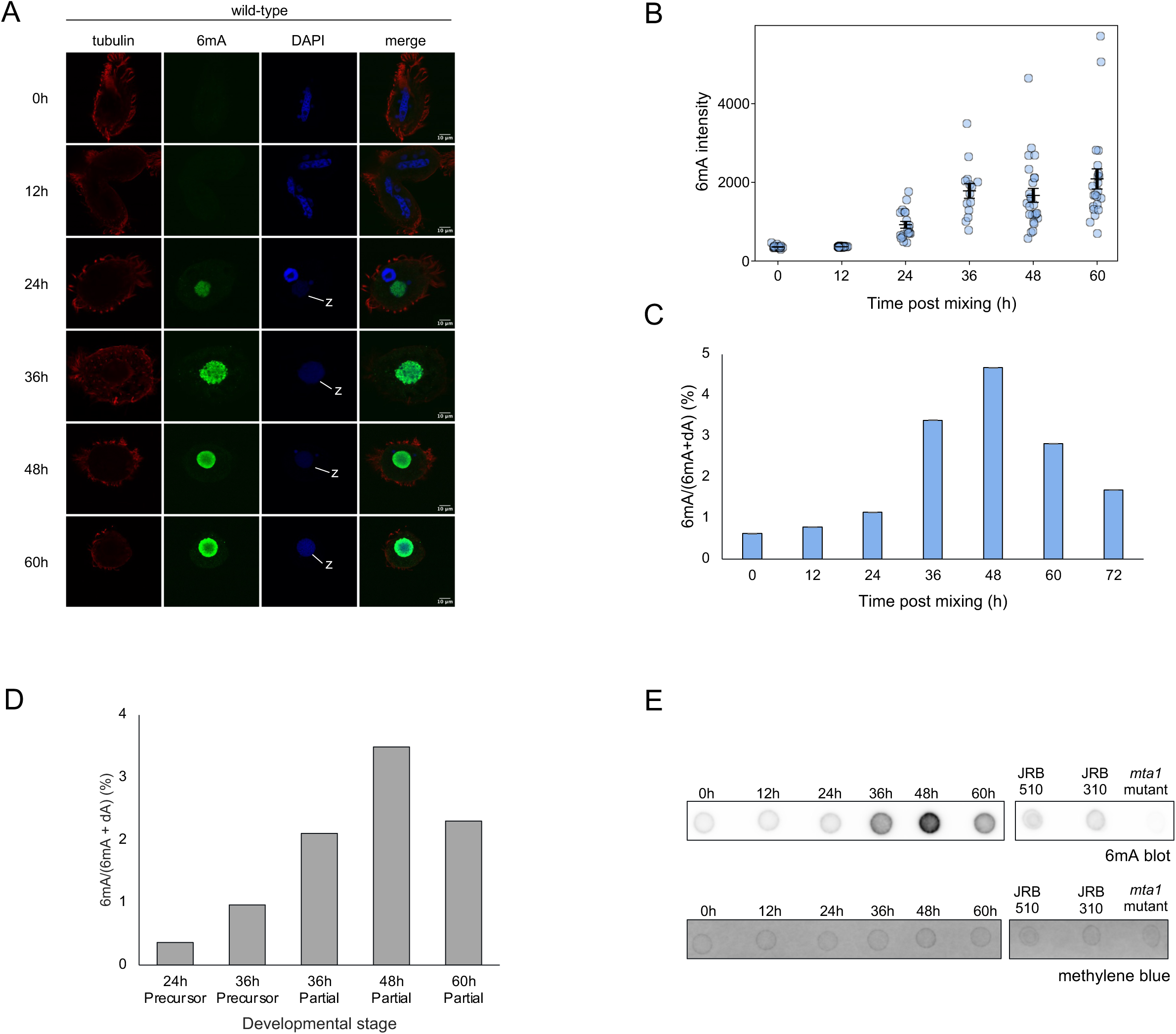
Adenine DNA methylation levels peak during development. **A.** Immunofluorescence analysis of 6mA (green) subcellular localization at indicated timepoints post-mixing of mating-compatible wild-type strains. DNA is labeled with DAPI (blue), and tubulin (red) denotes cell shape. Samples were treated with RNase. z: zygotic nucleus (anlage). **B.** Values of pixel intensities of the images shown in (A). Bars indicate mean ± standard error of the mean (SEM). Sample sizes were: 0h (n = 24), 12h (n = 43), 24h (n = 19), 36h (n = 14), 48h (n = 25), and 60h (n = 22). **C.** Quantification of 6mA abundance, relative to (dA+6mA) by mass spectrometry at the indicated timepoints during development. **D.** 6mA abundance, based on pooled SMRT-seq subreads, in either precursor or partially-rearranged reads (labeled “Partial”) at the indicated timepoints during development. Background of 6mA+dA is determined empirically by the number of A/T base-pairs in the genome with sufficient subread coverage (≥20) within each read set. 24h partially-rearranged reads were excluded as most appear to be somatic chromosome isoforms in the parental MAC that do not match the reference genomes. Precursor reads at later time points were also excluded as they become more likely to derive from non-developing micronuclei as rearrangement progresses. **E**. Dot blot analysis of 6mA during wild-type development (25ng total RNase-treated genomic DNA per dot) with methylene blue as a loading control.

To study 6mA enrichment at the sequence level, we performed PacBio single-molecule real-time sequencing (SMRT-seq)^37^ on total DNA isolated at 0, 24, 36, 48, and 60h of development^38,39^. To identify reads that most likely derive from the developing nucleus (anlage, plural anlagen), reads were mapped first to the reference somatic genomes of both mating-compatible *Oxytricha* strains (JRB310, JRB510)., Reads that perfectly mapped to either MAC reference genome were labeled “product” and binned separately, since they may derive from the macronucleus. The remaining reads were subsequently mapped to the available germline MIC reference genome for strain JRB310^23^ and then split into two categories of *perfectly-mapping* vs. *partially-mapping* (Figure S1B). At early–mid developmental timepoints (24–36h), reads that perfectly map to the MIC genome probably derive from unprocessed, precursor DNA in the polytenized anlagen^40^, while at mid–late developmental timepoints (36–60h), partially MIC-mapping reads likely represent partially-rearranged DNA in the developing nucleus. The first bin we call precursor reads, while the second bin we label partially-rearranged reads, reflecting their intermediate status. Both categories were included in further 6mA base-calling analysis. Partially-rearranged reads are most abundant at 36h–48h and remain detectable at 60h (Figure S1C and Table S1).

SMRT-seq analysis demonstrated increased 6mA levels across these developmental stages, compared to previously reported levels in vegetative, asexually growing cells^7^, with maximum levels observed in partially-rearranged 48h reads (Figure 1D). 6mA abundance was normalized to each dataset’s coverage of A/T base-pairs. Dot blot assays of total cellular DNA corroborated these dynamics (Figure 1E). Together, these results establish that 6mA specifically accumulates in the developing macronucleus and peaks at 48h post-mixing.

We next checked whether DNA methylation levels coincide with expression of the MTA1c methyltransferase catalytic subunit MTA1^7^ and its interacting partners. RNA-seq^41^ reveals increased MTA1 expression during development, consistent with our previous findings^7^, with a peak early in development (Figure S1D). While MTA9 expression remains low during development, we identified a paralog, MTA9-B (Contig17419.0, g11745)^25^, that is abundant and whose mRNA levels are similar to MTA1. In *Tetrahymena thermophila*, the MTA9-B ortholog AMT6 interacts with MTA1 (AMT1) during development^14^. This suggests that MTA9-B might serve as the developmental partner of MTA1 in *Oxytricha* instead of MTA9^7^. To test this, we generated a FLAG-tagged MTA1 transgenic line in an *mta1* mutant background (Figure S1E) and confirmed catalytic activity for 6mA addition in vegetative cells (Figure S1F). Immunoprecipitation using an anti-Flag antibody at 24h, coupled to mass spectrometry, demonstrated co-purification of FLAG-MTA1 with MTA9-B, but not MTA9 (Figure S1G), suggesting that MTA1 forms a developmental methyltransferase complex with MTA9-B to mediate 6mA deposition in the developing nucleus. Immunofluorescence analysis showed that FLAG-MTA1, which is expressed from the parental macronucleus, also localizes to anlagen at 24h post-mixing, supporting its proposed role in mediating 6mA deposition in development (Figure S1H).

### 6mA specifically marks retained DNA regions during development

Having established that both 6mA and MTA1 transcript levels increase during development, and that 6mA and MTA1 localize to the developing macronucleus, we next examined where 6mA is deposited during development. Pooled SMRT-seq subreads were used to identify methylation sites. Each 6mA position, and total A/T base-pairs sequenced with sufficient subread coverage (≥ 20) for 6mA calling, was assigned to one of six functional categories of the germline genome, defined in Feng *et al.* 2022^42^: either MDS (the only sequence category that is retained post-development), IES, TBE (Telomere-Bearing Element transposons^23^, Other Repeats^42^, Non-coding regions, or MIC genes. “MIC genes” represent a merged category of previously-annotated, predicted or confirmed germline-limited open reading frames (ORFs)^23,41^. This allowed inference of the methylation frequency within each genomic category (Figure 2A). While methylation levels are near the detection threshold among precursor germline genome-derived reads before the onset of development (0h post-mixing, Table S3), overall 6mA abundance increases dramatically during development, reaching 3.5% of total deoxyadenosines with sufficient subread coverage in 48h partially-rearranged molecules (Figures 1D and 2A). MDSs, in particular, appear to be the reservoir for most 6mA, climbing from 1.27% of total sequenced MDS A/T base-pairs in 24h precursor-mapped reads to almost 10% in 48h partially-rearranged reads (Figure 2A). At every developmental stage, 6mA levels are much higher within MDSs than any category of eliminated sequence, suggesting that 6mA deposition is specifically driven towards MDSs.

**Figure 2.**
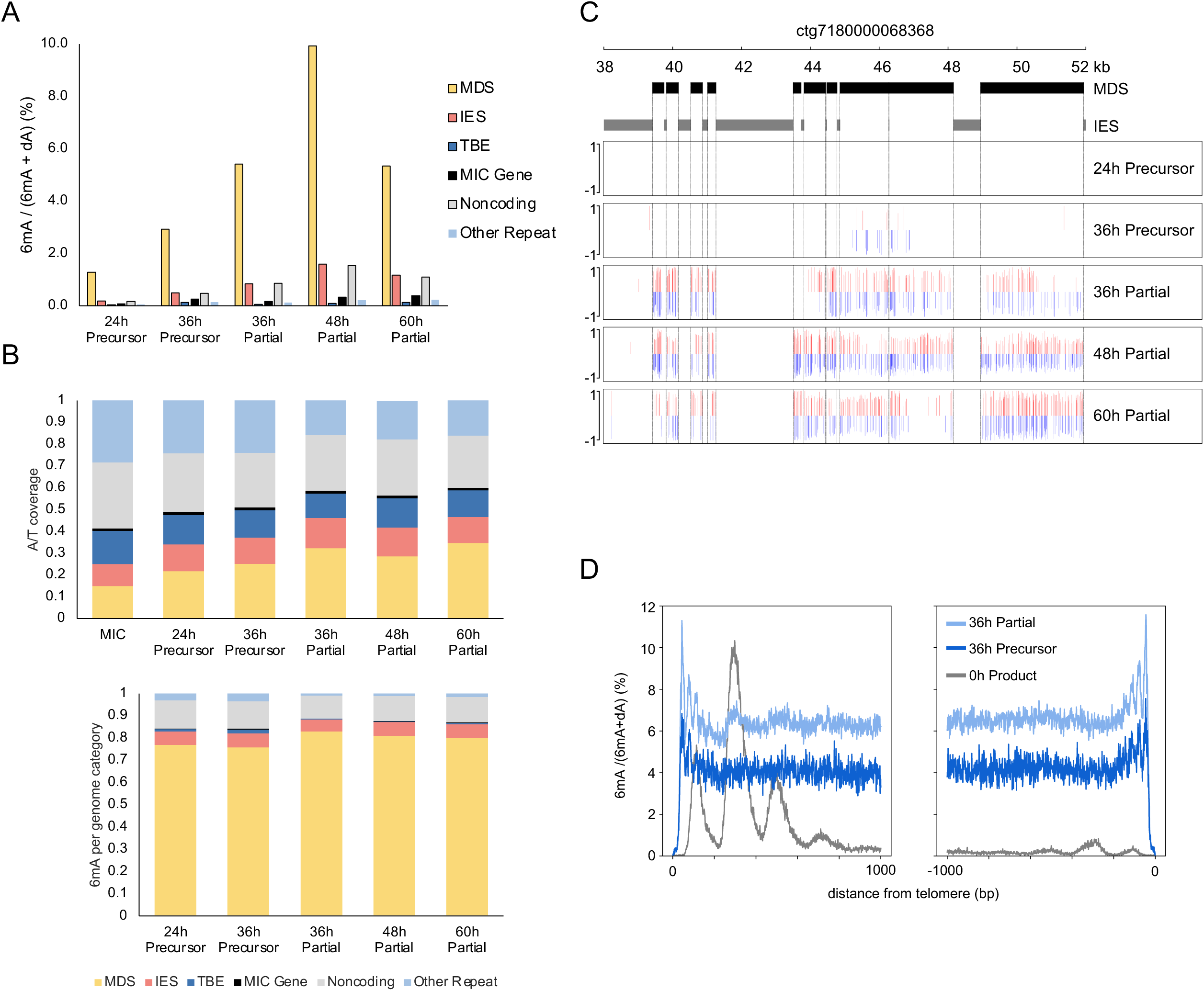
6mA specifically marks retained DNA regions during development. **A**. Percentage 6mA from pooled SMRT-seq subreads out of total sequenced A/T base-pairs, for the indicated developmental stages and germline genome categories (MDS, IES, Telomere-Bearing Element transposons^23^, Other Repeats, Non-coding regions, or MIC genes, as in ref.^42^). **B**. Proportion of genome coverage by SMRT-seq subreads (top panel) and of 6mA identified from pooled subreads (bottom panel) overlapping with each of six germline genome categories. Subread coverage exclusively counts A/T base-pairs. “MIC” refers to the proportion of A/T across genome categories in the reference germline genome^23^. **C**. 6mA methylation across an example 14kb germline genome region containing 10 MDSs (black boxes) and 11 IESs (gray boxes). 6mA calls detected from pooled SMRT-seq subreads are annotated on sense (red) and antisense (blue) DNA strands, with line height showing SMRT-seq subread methylfraction at each adenine (proportion of supporting subreads across all pooled data) (MDS, macronuclear-destined sequence; IESs, internal eliminated sequence). **D**. Meta-chromosome plots showing the proportion of 6mA [6mA/(6mA+dA)] in MDSs at 36h, relative to their MDS locations on somatic MAC chromosomes, for precursor (darker blue, 5,841 single-gene chromosomes) and partially-rearranged molecules (light blue, 19,124 single-gene chromosomes), plotting the first or last 1000 bp from telomere addition sites, excluding the telomere itself. The 6mA pattern in the parental macronucleus (0h post-mixing) is shown for reference (grey, 11,927 single-gene chromosomes). All chromosomes are aligned and oriented with their 5′ ends proximal to transcription start sites (TSSs).

We then measured the proportion of total 6mA that falls within each functional genomic category. While 6mA predominantly occurs within MDSs at all timepoints examined (Figure 2B, lower panel), MDSs comprise less than 15% of the germline genome (Figure 2B, upper panel). Though SMRT-seq reads become increasingly more enriched in MDSs, as other sequence classes are eliminated, MDSs never comprise more than 35% of total sequenced A/T base coverage. Nevertheless, over 75% of detected 6mA occurs within MDSs at every developmental stage, representing an enormous enrichment (2-tailed chi-square *p*-value < 2x10^-16^ for all stages). Precursor reads display 6mA enrichment on MDSs even at 24h, prior to the onset of DNA rearrangement. Germline-limited sequence categories are reciprocally depleted in 6mA, suggesting that 6mA may be a specific mark for retained DNA (Figures 2B and 2C).

While 0h SMRT-seq data (Figure 2D, gray) recapitulated our previous finding of promoter-proximal deposition of 6mA on MDSs in vegetative macronuclei^7^, a meta-chromosome analysis of MDSs destined for the first or last 1000 bases of over 5000 single-gene nanochromosomes displayed a strikingly different pattern of consistently high 6mA deposition during development (Figures 2D and S2A). Since the average length of single-gene nanochromosomes is just 2.7kb excluding telomeres, and many are <2kb ^27^, this suggests a contrastingly even distribution of 6mA across MDSs in the anlagen. This ubiquity is established as early as 24h in precursor reads, and persists through late development, even in 60h partially-rearranged reads and reaches its peak in 48h partially-rearranged molecules (Figure S2A). (The terminal 6mA spikes simply reflect a subtelomeric purine bias with 10bp periodicity^27,43^). These data establish that the more uniform pattern of 6mA deposition during development is completely different from the 0h or vegetative pattern associated with somatic MAC chromosomes and nucleosome positioning and suggest that 6mA may be a general or protective mark for MDSs during development.

Because pooled subread analysis aggregates signals from multiple molecules, it can artificially increase 6mA frequency estimates, especially in regions with high coverage. Therefore, to obtain a more precise estimate of 6mA abundance at the single-molecule level, we also quantified methylation within individual CCS reads. This single-molecule analysis confirmed pooled subread 6mA calling conclusions that abundance generally increases during development (Figure S2B), and that 6mA is highly enriched within MDSs (Figures S2C and S2D). Although overall 6mA levels are highest at 48h, single-molecule analysis revealed that 6mA abundance within MDSs is nearly as high in 36h precursor reads as in 48h partially-rearranged reads (Figure S2B). Furthermore, at 36h, the frequency of 6mA within MDSs is nearly twice as high among precursor reads as it is among partially-rearranged reads. Indeed, MDS 6mA frequency in 24h precursor reads is comparable to the level in 60h partially-rearranged reads (Figure S2B), consistent with a model of establishment of 6mA methylation on MDSs prior to DNA elimination and rearrangement.

We conclude developmental 6mA deposition occurs predominantly on MDSs, extends across the full length of MDSs, and pervasively targets MDSs, without the 5’ end bias or periodicity observed in asexual conditions. Importantly, 6mA becomes well established in the anlagen prior to DNA elimination. However, MDSs continue to be preferentially and increasingly methylated over the course of development.

### *mta1* mutant backcross disrupts DNA methylation during development

To test the developmental role of MTA1, we profiled 6mA dynamics in *mta1* X JRB510 backcrosses, which have developmental lethality and rarely survive past 60h^7^. Immunofluorescence revealed anlagen-localization of 6mA in *mta1* mutant backcrosses, similar to wild-type (Figure 3A). However, quantification showed significant reduction of nuclear 6mA in mutant backcrosses at 24h and 36h post-mixing but similar levels to wild-type by 48–60h (Figure 3B).

**Figure 3.**
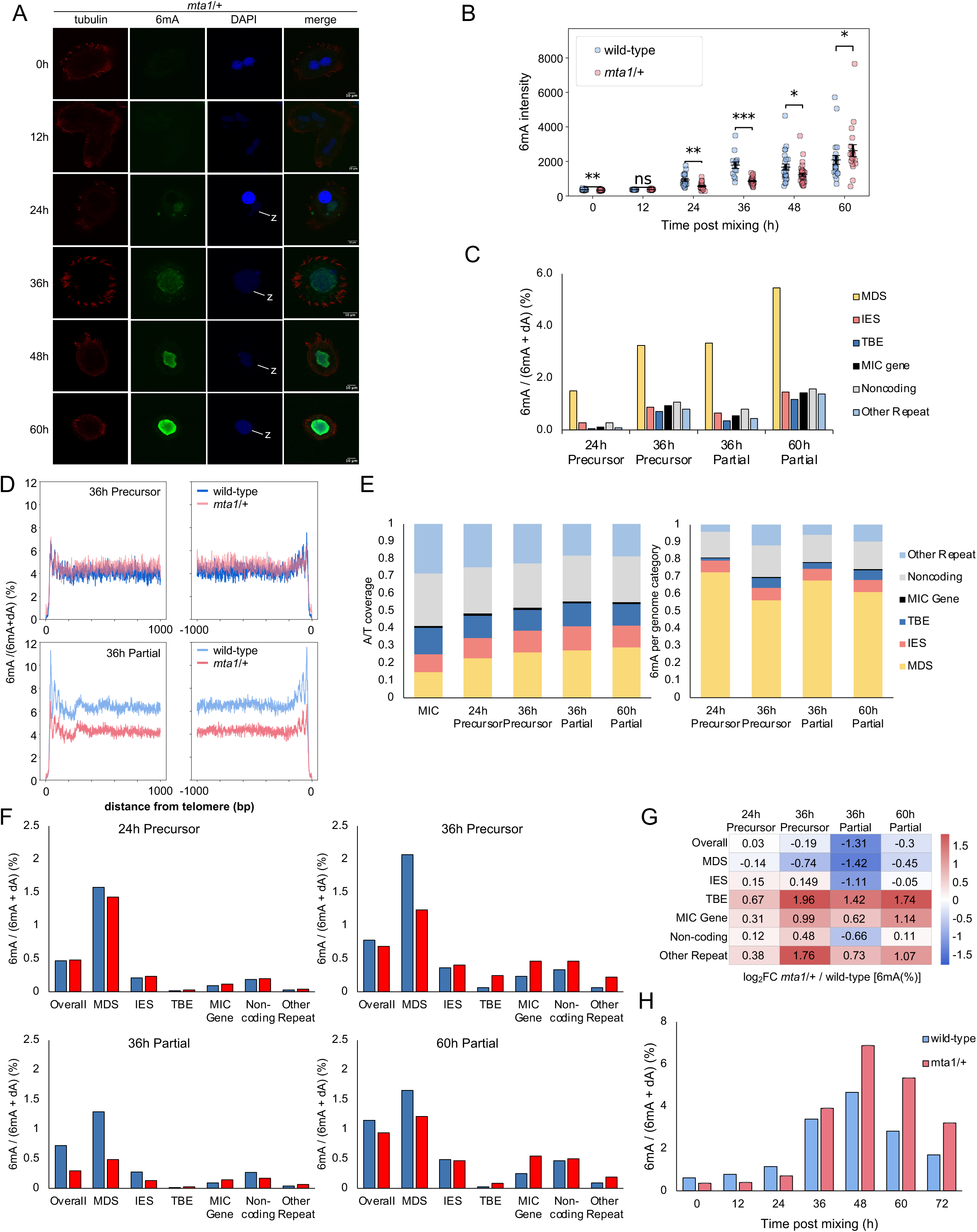
Developmental methylation is disrupted in an *mta1* mutant backcross. **A.** Immunofluorescence analysis of 6mA (green) subcellular localization in a backcross at indicated timepoints post-mixing of *mta1* mutant cells to JRB510 wild-type. DNA is labeled with DAPI (blue); tubulin (red) denotes cell shape. Samples were treated with RNase. z: zygotic nucleus (anlage). **B.** Values of pixel intensities of the images shown in (A). Bars indicate mean ± standard error of the mean (SEM). Stars indicate significance based on *p*-values from Mann-whitney U tests (* *p* < 0.05, ** *p* < 0.01 and *** *p* < 0.001). Sample sizes were the same as in Figure 1B for the wild-type and 0h (n = 27), 12h (n = 38), 24h (n = 19), 36h (n = 27), 48h (n = 34), and 60h (n = 19) for the *mta1* mutant backcross. **C.** Percentage 6mA from pooled SMRT-seq subreads out of total sequenced A/T base-pairs, for the indicated developmental stages and germline genome categories in the *mta1* backcross. **D.** Meta-chromosome plots as in Figure 2D, showing the proportion of 6mA [6mA/(6mA+dA)] in the developing 36h nucleus for precursor reads (top), contrasting wild-type (dark blue, 5,841 chromosomes) and *mta1* backcross (pink, 5,997 chromosomes) versus partially-mapping reads (bottom) for wild-type (light blue, 19,124 single-gene chromosomes) and *mta1* backcross (pink, 15,392 chromosomes). **E.** Proportion in the *mta1* backcross of genome coverage by SMRT-seq subreads (left panel) and of 6mA identified from pooled subreads (right panel) overlapping with each of six germline genome categories. Subread coverage exclusively counts A/T base-pairs. “MIC” (left) refers to the proportion of A/T across genome categories in the reference germline genome^23^. **F.** Percentage 6mA on individual SMRT-seq CCS (circular consensus sequencing) reads in wild-type and *mta1* backcross out of total sequenced A/T base-pairs, at the indicated developmental stages and germline genome categories. **G.** Log_2_ fold-change (log_2_FC) of the data in Figure 3F, comparing *mta1* backcross vs. wild-type cells at the indicated developmental stages and genomic categories, called from individual CCS reads. Log_2_FC > 0 (red) indicates higher 6mA abundance in *mta1* backcross and log_2_FC < 0 (blue) indicates higher 6mA abundance in wild-type. **H.** Quantification of 6mA abundance, relative to (dA+6mA) by mass spectrometry at the indicated timepoints during development, in wild-type (JRB310, blue) or *mta1* mutant (red) cells mated to JRB510.

To quantify 6mA abundance and distribution in *mta1* mutant backcrosses, we performed SMRT-seq on 3 developmental timepoints. Anlagen-derived reads were categorized as before (Figures S1B and S3A). As expected, parental 6mA at the onset of conjugation is approximately half as abundant in the mutant backcross as in wild-type mating, with a similar distribution (Figure S3B). Pooled subread 6mA calling showed a redistribution of 6mA in the mutant backcross, with 39% less 6mA on MDSs mid-development, among 36h partially-rearranged reads, and more 6mA on germline-limited categories, especially TBEs and other repeats (Figures 2A, 3C)^42^. Meta-chromosome analysis, as above, showed that this decrease occurs relatively evenly across MDSs, compared to wild-type, and is only detectable on partially-rearranged DNA but absent from the precursor stage at 36h of development (Figure 3D). We did not detect a similar loss of 6mA at very early (24h) or late (60h) timepoints in the pooled-subread analysis (Figure S3C). Although the majority of 6mA is still on MDSs in the mutant backcross, this enrichment is substantially less than in wild-type (Figure 3E). Rather, 6mA is consistently less depleted in germline-limited sequences in the *mta1* mutant backcross (Figure S3D), suggesting less accurate targeting of the modification to retained sequences.

To more precisely quantify the frequency of methylation in the mutant backcross, we called 6mA on individual CCS reads, as before. This single-molecule analysis revealed that methylation frequency within MDSs decreases in the *mta1* mutant backcrosses in anlagen-derived DNA from all developmental stages examined (Figure 3F), except the 24h precursor. Individual CCS analysis further demonstrated higher 6mA levels in germline-limited DNA categories, on the other hand, in the mutant backcross, especially repetitive elements. 6mA was most elevated on germline-limited regions in 36h precursor molecules and 60h partially-rearranged molecules, indicating continued aberrant methylation. 24h precursor molecules exhibited the least difference between wild-type and mutant backcross, suggesting that either the reduced dosage is sufficient or that another methyltransferase might contribute to initial *de novo* 6mA deposition (Figures 3F and 3G). This is consistent with our previous demonstration that the MTA1 ortholog in *Tetrahymena* prefers a hemimethylated substrate *in vitro*, though it is also a *de novo* methyltransferase^7^. Germline-limited 6mA levels on single molecules in the mutant backcross were generally still lower than would be expected if randomly deposited, but MDS enrichment was even more reduced than in pooled subread analysis (Figures S3E and S3F).

LC-MS/MS [6mA/(6mA+dA)] and 6mA dot blot analysis provided complementary, whole-cell measurements. At early timepoints (12–24h), the mutant backcross exhibits lower global 6mA than wild-type, in accordance with immunofluorescence and SMRT-seq data (Figures 3H, S3G and S3H). As development proceeds, these values converge, and by late rearrangement, whole-cell mutant backcross 6mA levels exceed wild-type. Because both LC-MS/MS and dot blot assays are nucleus-agnostic, these late increases likely reflect stalled processing of methylated DNA in the mutant backcross (e.g., the persistence of partially-processed molecules) or developmental asynchrony. Diminished MTA1 gene dosage therefore perturbs both where and when 6mA is deposited, not the cell’s overall capacity to methylate DNA. We conclude that the *mta1* mutant backcross affects timing and location of 6mA deposition and that wild-type levels of MTA1 are required to target 6mA to MDSs during development, and to prevent aberrant deposition on germline-limited sequences, ensuring faithful methylation patterns during post-zygotic development.

### 6mA deposition is less precise in the mta1 mutant backcross

We previously reported that MTA1 preferentially methylates ApT motifs in asexually growing *Oxytricha*^7^. To investigate the motif specificity of 6mA in *Oxytricha* development, we examined the dinucleotide context of 6mA in individual CCS reads. While the vast majority of 6mA events in the mutant backcross still occur at 5’-ApT-3’ motifs, as in vegetative cells^7^, there is proportionally more hemi-methylated ApT and methylation on non-ApT motifs in the backcross, especially in partially-rearranged molecules (Figure 4A), again consistent with the possibility that *Oxytricha* MTA1 may be a better maintenance than *de novo* methyltransferase, like its *Tetrahymena* ortholog^7,44^. The mutant backcross has notably less hemi- and fully-methylated ApT dinucleotides in partially-rearranged reads, especially mid-development, 36h post-mixing (Figure 4B), and the distance between two adjacent 6mA sites on individual DNA molecules is also greater at all developmental stages (Figure 4C). Intriguingly, the vast majority of neighboring 6mA events in wild-type occur within 27bp of each other—the length of *Oxytricha* piRNAs that also mark MDSs^31,32^. This raises the intriguing possibility that a single piRNA can guide a unique 6mA addition event.

**Figure 4.**
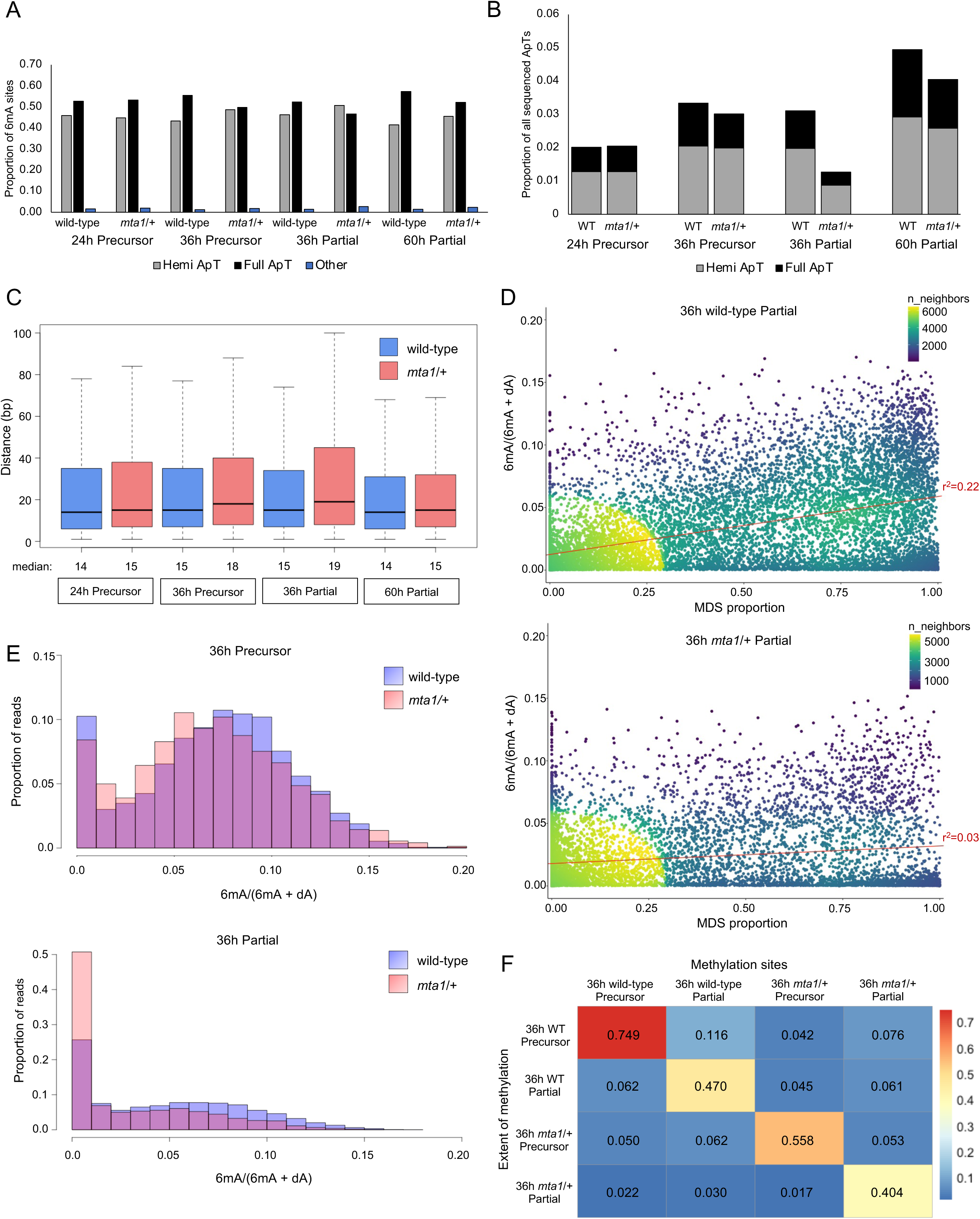
Mutation of *mta1* leads to imprecise 6mA deposition in backcross. **A.** Distribution of 6mA across hemi-methylated ApT (grey), fully-methylated ApT (black), and non-ApT sites (“Other”, blue), measured by SMRT-seq at single molecule resolution, at the indicated developmental stages. Samples derive from either a wild-type cross or an *mta1* mutant backcross (*mta1* / +). Values represent the proportion of all 6mA-containing loci per category. **B.** Proportion of sequenced ApT motifs that are either hemi-methylated (grey) or fully-methylated (black) at single molecule resolution. Values reflect the fraction of all ApT motifs in CCS reads above quality thresholds (≧20 subreads). As in other plots, developmental stages (precursor or partially-rearranged) and matched timepoints are shown for both wild-type (JRB310) and *mta1* backcrossed to JRB510 wild-type. WT: wild-type. **C.** Boxplot showing the distribution of distances between neighboring 6mA sites within single CCS reads, excluding the 1 bp distance between 6mA on dimethylated ApT motifs. Developmental stages (precursor or partially-rearranged) and timepoints are shown for wild-type or *mta1* crossed to JRB510. **D.** Proportional MDS content of each CCS read plotted against the extent of 6mA methylation detected on that read, for 36h partially-rearranged CCS reads from wild-type and *mta1* backcross SMRT-seq datasets. Values reflect reads passing quality thresholds (≧ 20 subreads) and containing at least one 6mA site. Dot color represents density of data. The red line indicates the simple linear regression (*r*^2^ is the square of the associated correlation coefficient). **E.** Histograms of 6mA abundance across 36h CCS reads, shown for precursor reads (top) and partially-rearranged reads (bottom) in wild-type (blue) and *mta1* backcross (red) backgrounds, with overlap in purple. 6mA abundance is calculated per read as 6mA divided by all sequenced adenines in that read. The y-axis indicates the proportion of reads in each 6mA-abundance bin. Only reads with high MDS content (≥50% MDS) and containing at least one 6mA site are included. **F.** Site-specific extent of methylation at 36h at single molecule resolution for each developmental stage and genetic cross. For the CCS reads indicated in each row, the extent of methylation is measured at genomic positions where 6mA was found in the CCS reads in each column. Values within the heatmap represent total 6mA calls divided by total CCS read coverage at the corresponding 6mA sites. (Sites with no CCS coverage in a given read category do not affect calculation.) Entries along the diagonal measure penetrance of 6mA in the given set of CCS reads. CCS reads with fewer than 20 subreads are excluded.

To study the ubiquity of 6mA on MDSs, we compared the 6mA abundance on each individual CCS read at 36h to the MDS proportion on that molecule (Figures 4D and S4A). While some reads devoid of 6mA may be undergoing endoreplication, methylated wild-type reads displayed a roughly linear relationship between MDS density and 6mA abundance, especially in precursor reads. In the *mta1* mutant backcross, however, this correlation was both greatly reduced in precursor reads and entirely absent from partially-rearranged reads, suggesting global disruption of 6mA deposition. On partially-processed molecules with no MDSs, the average 6mA in the mutant backcross (0.36%) is nearly twice that of wild-type (0.21%), while the converse is true for majority-MDS reads (2.9% in mutant backcross vs. 4.9% wild-type) (Figure 4D). On those MDS-rich reads, the mutant backcross displays generally less methylation on precursor DNA (Figure 4E), with 6mA less evenly present on both DNA strands, relative to wild-type, consistent with less methyltransferase activity (Figure S4B). Partially-rearranged DNA in the mutant backcross contains more reads that are nearly completely unmethylated (Figures 4E, S4C), suggesting global disruption of DNA methylation.

With single nucleotide resolution, we then checked the frequency of recurrence of methylated sites at specific developmental stages. We extracted genomic positions of 6mA calls in each 36h read category and defined penetrance as the proportion of mapped CCS reads showing methylation at each site. For example, among all discrete 6mA sites in 36h wild-type precursors, 74.9% of precursor reads covering those sites are methylated. While partially-rearranged reads display more unique methylated sites, likely a consequence of more CCS reads, penetrance at 6mA sites is highest in wild-type precursor reads (Figure 4F and Table S1). Penetrance is also always higher in wild-type than mutant backcross at each developmental stage. When we compared 6mA positions across genotypes and developmental stages, we found that sites methylated in one read category are rarely recurrently methylated in another, especially in the mutant backcross. Wild-type methylation levels on precursor molecules at sites also methylated in partially-rearranged molecules (11.6%) are more than twice as frequent in wild-type than in *mta1* mutant backcrosses (5.3%), indicating less robust methylation targeting in the mutant. When considering specific bases methylated at 36h in wild-type, methylation frequency falls dramatically in the mutant backcross (Figure S4D and S4E). These results suggest that *mta1* mutation leads to more sporadic 6mA methylation in *Oxytricha*, both improperly timed and often imprecise at single nucleotide resolution.

### *mta1* mutant backcross exhibits a developmental delay

To determine whether reduced MTA1 gene dosage in mutant backcrosses impacts timing of DNA rearrangements, we introduced a developmental index that allows us to quantify junction processing as the number of processed (i.e., MAC-like) junctions divided by the total number of SMRT-seq reads spanning those junctions in wild-type vs. *mta1* mutant backcross datasets. Because consecutively-numbered MDSs in *Oxytricha* can be out of order (scrambled) or inverted in the precursor genome, we are unable to use the IES-retention score that is used for species with only IES elimination^45^. By analyzing an intersection of over 65,000 pointers with representation in both wild-type and mutant backcross datasets at 36 and 60h of development, we find that wild-type crosses display a greater proportion of fully processed junctions (Figure 5A). By contrast, we did not observe a difference in non-MAC/non-MIC (i.e., “Other”) pointer categories аt 36h post-mixing, consistent with a developmental delay rather than large errors such as incorrect junction formation (Figure S5A).

**Figure 5.**
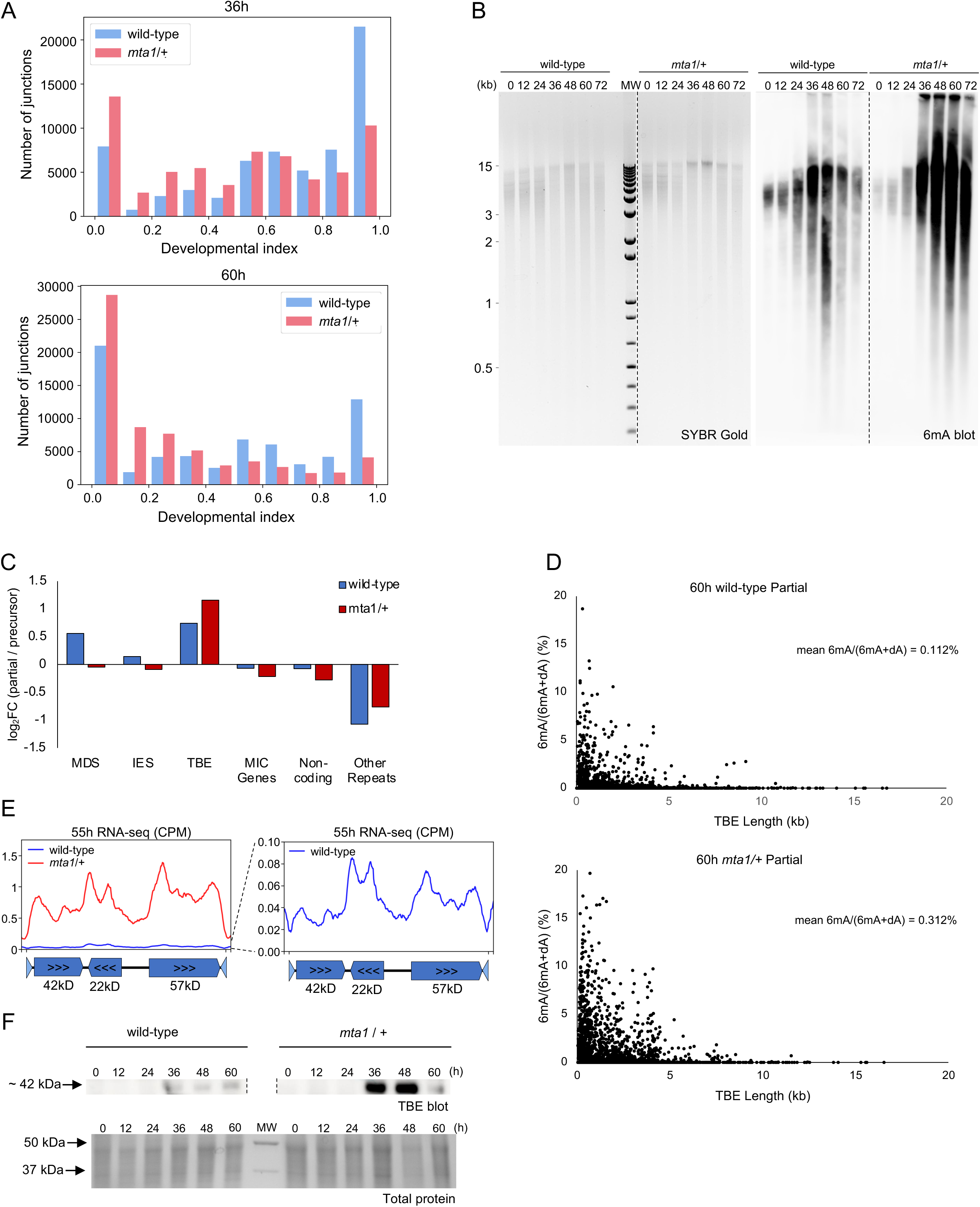
*mta1* mutant backcross exhibits a developmental delay. **A.** Developmental index (proportion of CCS reads covering fully processed junctions for a given junction) shown for the shared set of well-covered junctions at 36h (67,049 junctions) and 60h (63,911 junctions) among partially-rearranged molecules in wild-type (blue) or *mta1* backcross backgrounds (pink). **B.** Immuno-Southern hybridization of total genomic DNA probed with anti-6mA antibody (right) with SYBR Gold loading control (left) for wild-type and *mta1* backcrosses at the indicated timepoints post-mixing. **C.** Log_2_ fold-change (log2FC) in A/T coverage across the six germline genome categories at 36h, comparing partially-rearranged vs. precursor CCS SMRT-seq reads, in wild-type (blue) and *mta1* backcross (red) backgrounds. **D**. 6mA abundance on individual TBE transposons of varying lengths at 60h, measured by single-molecule 6mA calling. Each TBE shown was required to have at least 100 A/T base-pairs of coverage by CCS reads. Outlier TBEs with >10X coverage were excluded because of potential false mapping, as were TBEs annotated as longer than 18kb. In total, 9462 TBEs in wild-type and 8826 in *mta1* backcross passed these filtering criteria. The mean 6mA/(6mA+dA) for all included TBEs was 0.112% in WT and 0.312% in the *mta1* mutant backcross. For TBEs of canonical length (3–5kb), mean 6mA abundance was 0.035% in wild-type and 0.143% in the mutant backcross. **E.** Average counts-per-million-reads (CPM) of length-normalized “ideal” TBEs from 55h RNA-seq data, averaged across three technical replicate libraries. Seventy-nine TBEs were classified as “ideal” on the basis of canonical number, length, and orientation of their three ORFs, as well as overall length of the transposon. Left panel: TBE expression profile from wild-type (blue) and *mta1* mutant backcross (red). Right panel: expanded view of the blue wild-type profile. Both datasets show three sets of peaks, corresponding to transcription start and end of the three transposon ORFs, shown in the schematic TBE model below. **F.** Western blot detection of TBE transposase in wild-type and *mta1* backcross at the indicated time-points of development using a TBE transposase-specific antibody^47^. Below, total protein staining is shown as a loading control.

Additional evidence supporting impaired DNA processing is the accumulation of high molecular weight DNA in *mta1* mutant backcrosses, as indicated by immuno-Southern hybridization of DNA to an anti-N⁶-methyladenine antibody (Figure 5B). We note that previous experiments also demonstrated accumulation of high molecular weight DNA when TBE transposase levels were reduced by RNAi, consistent with stalled genome rearrangement^38^. Relative to precursor molecules, 36h partially-rearranged SMRT-seq reads in the *mta1* mutant backcross are also less enriched in MDSs and exhibit greater retention of TBEs and other repeats, consistent with developmental delay (Figure 5C).

Impairment of DNA processing is further supported by changes in transcription: RNA-seq reads from wild-type and *mta1* mutant backcrossed cells at 48h, when 6mA peaks, and 55h, before most cells die in the mutant backcross^7^ display less mapping to the product MAC genome in the mutant backcross (Figure S5B). Reads derived from TBEs, whose encoded proteins are only expressed during development^38^, and which are more methylated overall in the mutant backcross (Figure 3G), are more abundant in the mutant backcross RNA-seq (Figure S5C). The methylation status of individual TBEs is also broadly and consistently more methylated in 60h partially-rearranged reads in *mta1* mutant backcrosses, based on analysis of individual CCS reads that map to >8,000 well-covered TBEs (Figure 5D). Detailed analysis of 79 complete TBE transposons with 3 intact reading frames^46^ showed that transcription is generally enriched across the locations of the 3 ORFs in mutant backcrossed cells (Figure 5E). These same 79 complete TBEs exhibit threefold higher 6mA levels in the 60h partially-rearranged reads in the mutant backcross, with comparable methylation frequency in coding and linker regions (Figure S5D). Upregulation of TBE RNA abundance in the mutant backcross was not associated with specific upregulation of other germline-limited ORFs (Figure S5C), suggesting either specific TBE derepression or, we postulate, a weaker ability to shut down TBE transcription if the transposons are protected against DNA deletion by increased 6mA methylation. To test whether changes in TBE transcription were accompanied by altered transposase protein levels, we used a TBE2.1 antibody^47^ in western hybridization and found that transposase signal increased in mutant backcrosses at 48-60h (Figure 5F). This transposase dysregulation may contribute to the mutant phenotype. Together, these data demonstrate that the *mta1* mutant backcross has a significant developmental delay in genome rearrangements, including delayed junction processing, reduced transcription of mature, somatic genes in the new MAC, and altered transposon expression and elimination.

### 6mA is a PIWI protein-dependent, epigenetic mark for DNA retention

piRNAs mark genomic regions for retention in *Oxytricha*^31^ but appear much earlier than genome rearrangements. Since 6mA peaks later than piRNAs in development and colocalizes with the *Oxytricha* PIWI subfamily protein Otiwi1 in the anlagen at 24h (Figure 1A), we hypothesized that 6mA may function downstream of piRNAs as a secondary mark for MDS retention. Previously, we showed that piRNA production requires Otiwi1. To test whether 6mA deposition also requires Otiwi1 function, we examined 6mA levels in a cross between two *otiwi1* mutant lines containing insertions that disrupted the Otiwi1 reading frame^31^. Sequencing confirmed gene disruption via IES retention, and immunofluorescence demonstrated the absence of expression of the Otiwi1 protein (Figure S6A), confirming loss-of-function lines. We then assessed 6mA levels during development in the progeny of this cross between mating-compatible *otiwi1* mutant strains. While immunofluorescence revealed strong nuclear 6mA in wild-type control cells at 24h, 6mA signal was markedly reduced in *otiwi1* mutants (Figure 6A). Quantification showed a significant decrease in nuclear 6mA intensity (*p* = 0.01, n ≥ 3, *t*-test) (Figure 6B) and SMRT-seq confirmed nearly eight-fold reduction of 6mA base calls in *otiwi1* mutant 24h precursor reads (Figure 6C). DNA immuno-Southern hybridization also confirmed reduced 6mA accumulation in the *otiwi1* mutant background (Figure 6D). Together, these data indicate that Otiwi1, and by extension, the piRNA pathway, are required for 6mA marking during development.

**Figure 6.**
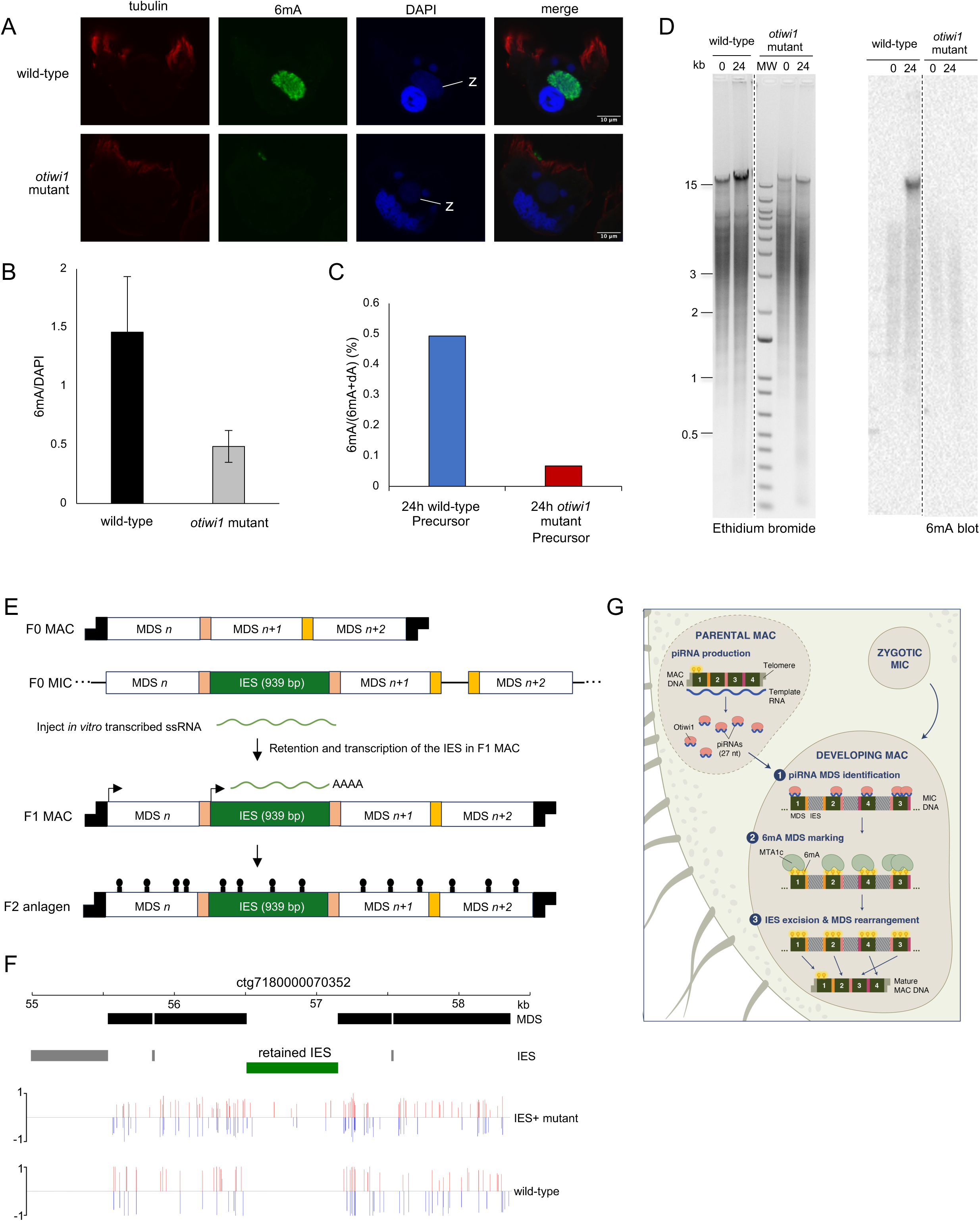
6mA is a PIWI protein-dependent, epigenetic mark for DNA retention. **A.** Immunofluorescence of 6mA (green) subcellular localization, in wild-type and *otiwi1* x *otiwi1* mutant crosses at 24h post-mixing. DNA is stained with DAPI (blue), and tubulin (red) outlines the cell shape. Z: zygotic nucleus (anlage). **B.** Average values of integrated pixel intensities of the images shown in (A), normalized relative to DAPI. Error bars represent standard deviation; *p* = 0.01, *t*-test, *n* ≧ 3. **C.** Percentage of 6mA detected in pooled SMRT-seq subreads, divided by total sequenced A/T base-pairs covered by ≥20 subreads, in 24h precursor reads from wild-type control and *otiwi1* x *otiwi1* mutant crosses. The *otiwi1* mutant shows a global reduction in 6mA abundance. **D.** Immuno-Southern analysis of total genomic DNA probed with anti-N⁶-methyladenine antibody (right) with corresponding Ethidium Bromide loading control (left) in wild-type and *otiwi1* x *otiwi1* mutant crosses at 0 or 24 hours post-mixing. The reduced signal in the PIWI mutant suggests reduced global 6mA levels. **E.** Schematic of transgenic line (misexpression mutant) with programmed retention of a transcribed germline-limited gene.^41^ Synthetic single-stranded RNA (ssRNA) covering an IES that contains a germline-limited gene was microinjected into developing cells to program long IES retention. This leads to the formation in the F1 progeny of a telomere-capped nanochromosome that retains the MIC-limited gene in the retained IES (green), together with surrounding MDSs (white). The retained IES is now transcribed during vegetative growth^41^. During the next sexual cycle, the retained IES, effectively a *de novo* MDS, can be a source of new piRNAs, as well as 6mA deposition on the IES in the F2 anlagen, leading to observed transgenerational inheritance of the retained IES. **F.** 6mA methylation tracks across a 3.5 kb germline region containing 4 MDSs (black boxes) and 3 IESs (gray boxes, one long and two short), plus the retained IES (green box) in a backcross between JRB510 and the IES+ line in panel E. 6mA calls detected from pooled SMRT-seq subreads are recovered on both sense (red) and antisense (blue) DNA strands, with line height showing SMRT-seq subread methylfraction at each adenine (proportion of supporting subreads across all pooled data). **G.** Schematic model for RNA-guided deposition of 6mA on MDSs leading to their protection against DNA deletion during genome rearrangement. A copy of the zygotic MIC differentiates to form the new MAC, and parentally-inherited piRNAs mark MDSs which leads to 6mA DNA methylation, which then confers protection against DNA deletion during assembly of new macronuclear chromosomes.

To test whether RNA-guided DNA retention is sufficient to program 6mA deposition on germline-limited DNA, we first examined a modest 62bp retained IES whose retention had been programmed by RNA injection to construct the *mta1* mutants^7^. On this short IES, we identified two instances in an *mta1* mutant backcross of 6mA deposition on partially-rearranged 36h and 60h SMRT-seq reads (Figure S6B). Reads from the wild-type cross had no 6mA within this IES. Given the small length of this IES, we analyzed a mutant *Oxytricha* line in which a 939bp IES that contains a germline-limited gene was artificially protected against deletion by microinjection of a synthetic RNA covering the IES^41^. This line produces a mature nanochromosome with the retained IES as a 939bp insertion (Figure 6E). SMRT-seq detected the emergence of 6mA at 36h across the retained IES on both strands (Figure 6F), despite a total absence of 6mA on this region in the wild-type, demonstrating that RNA–guided IES retention leads to *de novo* 6mA deposition on the retained DNA sequence.

Together, these experiments support a model in which 6mA acts as a functional, Otiwi1-dependent epigenetic mark that protects genomic regions against DNA deletion. Additionally, these observations and experiments demonstrate a link between RNA-guided sequence recognition and DNA methylation *via* the MTA1 complex to facilitate accurate genome remodeling during development.

## Discussion

Here we identify DNA N6-methyladenine (6mA) as a key epigenetic mark deposited on retained DNA segments during genome editing and reorganization in *Oxytricha* development. We find that 6mA accumulates in developing nuclei, before DNA elimination, and reaches its maximum levels 48-60 hours post-mating, especially when large-scale chromosome rearrangements take place. Furthermore, the 6mA marks are concentrated and relatively evenly distributed on the macronuclear-destined sequences (MDSs) that resist DNA deletion. By 48h, nearly 10% of deoxyadenines in MDS sequences can be methylated, while most eliminated DNA remains unmethylated. Development in an *mta1* mutant background disrupts this enrichment, leading to reduced 6mA presence on MDSs and increased 6mA on germline-limited sequences. Loss of the PIWI protein, Otiwi1, completely abolishes the detection of 6mA in the developing macronucleus, suggesting an upstream role for the piRNA pathway in 6mA deposition on retained genome segments.

This discovery of an interplay between the piRNA pathway and 6mA DNA methylation during *Oxytricha* development solves a longstanding paradox in the field. Otiwi1-bound 27-nt piRNAs accumulate on MDSs early in development but peak prior to genome rearrangements and then vanish before chromosome maturation^31^. This raises the question of how piRNAs transfer the epigenetic memory for DNA protection. The present study shows that 6mA marks supply this epigenetic memory. Artificially programming of an IES for retention induces *de novo* 6mA addition, demonstrating that retention alone is sufficient to trigger DNA methylation in the next generation. Together these results suggest that piRNAs recruit, directly or indirectly, the MTA1 complex to deposit 6mA on retained DNA (Figure 6G). While loss of Otiwi1 eliminates both piRNAs^31^ and 6mA marking, leading to early developmental arrest, *mta1* backcrosses can still initiate rearrangement, but arrest later. This also supports a model in which Otiwi1–piRNA complexes provide sequence-specific targeting of MDSs, and then MTA1 installs 6mA as a protective epigenetic mark. Long, non-coding, template RNAs^29^ can then have the opportunity to program MDS linkage and reordering, together with transposase-mediated DNA cleavage^38,47^ of unprotected germline-limited sequences. While our findings suggest a connection between the piRNA pathway and 6mA methylation during development, we did not find evidence of a direct interaction between the MTA1c complex and Otiwi1. However, in our previous study that identified Otiwi1^31^, immunoprecipitation with an anti-Piwi1L antibody followed by mass spectrometry also identified two peptides corresponding to MTA9-B, which here we identified is part of the MTA1c methyltransferase complex during development. This hints toward a functional interaction between DNA methylation and Otiwi1.

This partnership between small RNAs and DNA methylation echoes a broader theme of RNA-guided DNA methylation across eukaryotes. Two key precedents for small RNA-guided DNA methylation are siRNA-directed 5-methylcytosine (5mC) DNA methylation in plants^48^ and piRNA-directed 5mC DNA methylation in mammals^49,50^. Note that these cytosine methylation pathways typically target and silence transposons in the germline and can also direct histone methylation^48,51,52^. While RNA-guided chromatin regulation is an ancient principle, *Oxytricha* provides an orthogonal epigenetic strategy, where piRNAs and 6mA supply an activating mark, preserving, rather than silencing DNA. This may also convert a transient piRNA signal into a stable, protective mark for MDSs. The methylation of adenine further illustrates the plasticity of RNA-guided DNA methylation pathways.

In other ciliates, small RNAs and chromatin modifications more often target germline-limited DNA for elimination^53–60^. For example, in *Tetrahymena*, scnRNAs guide H3K9 methylation to IESs^61–68^, which are much longer in *Tetrahymena* than *Oxytricha*, and there is no evidence that 6mA guides DNA rearrangements in *Tetrahymena*^69^. Additional chromatin marks may also contribute in *Oxytricha*, in which 5mC accumulates in the degrading parental macronucleus^30^. Whether 6mA interacts with specific pathways remains unknown, but it does reduce nucleosome occupancy^7–14^, potentially exposing chromatin. Future studies exploring such interplay will be key to understanding the epigenetic choreography of genome remodeling. The roles of 6mA in ciliates can also vary further: disrupting *Paramecium*’s only DNA adenine 6mA methyltransferase did not detectably alter genome rearrangement^70,71^. The distantly-related ciliate *Loxodes magnus* lacks extensive genome rearrangement entirely but uses 6mA to differentiate its expressed MAC from the archival MIC genomes^20^, while *Pseudocohnilembus persalinus* shows reduced 6mA during encystment^72^.

The MTA1 complex is essential for both placing and propagating 6mA on MDSs. In *mta1* mutant backcrosses, methylation declines on MDSs, while germline-limited sequences acquire ectopic 6mA. This dysregulation presumably leads to a failure to complete the precise nuclear developmental program and eventual cell death, as *Oxytricha* development requires rearrangement of over 200,000 MDSs to create functional genes^23^. This underscores the importance of selectively targeting 6mA to somatic-destined DNA. TBE transposons, on the other hand, pervasively gain 6mA across thousands of copies in the developing genome, with RNA-seq and western analysis supporting concomitant increase in TBE-derived transcription in mutant backcrosses. 6mA acquisition on transposons could promote TBE expression indirectly, by blocking their excision rather than by directly activating transcription. Intriguingly, perhaps related to their expression, TBE chromatin becomes more accessible during wild-type development^73^, which offers a simple explanation for its increased methylation in the *mta1* mutant backcross. TBE dysregulation may be a primary cause of the developmental delay in the *mta1* backcross, given the observed excess of unprocessed junctions and accumulation of high molecular weight DNA.

The remodeling of 6mA during post-zygotic development in *Oxytricha* comes with an elegant twist: it may not require the presence of an active 6mA eraser, because endoreplication during late development would erase the developmental 6mA pattern while simultaneously increasing the ploidy of MAC nanochromosomes to an average copy number of approximately 1,900^22,27^. Our identification in the present study of 6mA as an MDS-specific, piRNA pathway-dependent mark that protects DNA during development therefore provides the key, missing epigenetic link in RNA-guided genome editing in *Oxytricha.* These findings also underscore how DNA modifications can direct complex developmental programs, and raise the question of whether similar strategies may exist in other organisms. More broadly, this work illustrates the power of noncanonical model systems like *Oxytricha* to expand our fundamental knowledge of epigenetic mechanisms that regulate and modify genome architecture and function.

## Supplemental Figures

**Figure S1. Adenine DNA methylation levels peak during development, related to Figure 1. A.** MRM chromatograms (Multiple Reaction Monitoring chromatogram) generated by Liquid Chromatography-Tandem Mass Spectrometry (LC-MS/MS) of N-6-methyladenosine (6mA, here labeled as dmA) at different timepoints; X-axis: retention time; Y-axis: mass response. **B**. Schematic representation of the bioinformatic pipeline used to bin PacBio CCS reads. Precursor (24-36h) and partially-rearranged (36-60h) reads derived from the developing somatic nucleus were used for analyses. JRB310 and JRB510 are the reference parental strains used in wild-type matings; MAC: macronucleus; MIC: micronucleus. **C.** Proportion of total SMRT-seq CCS reads assigned to each read bin (product, precursor, partially-rearranged and unmapped) at the indicated developmental timepoints in hours (see Table S1 for read counts). **D**. Developmental transcript abundance in RNA-seq of MTA1, MTA9, and MTA9-B at the indicated timepoints post-mixing in hours (h); TPM values (Transcripts Per Million) calculated from published RNA-seq datasets^41^. Gene identifiers used for quantification were: MTA1 (Contig12701.0.0; g10712), MTA9-B (Contig17419.0; g11745), and MTA9 (Contig1237.1; g24932) from the *Oxytricha trifallax* MAC genome^27^. Error bars show standard deviation across replicates. **E.** Top panel: Schematic of the FLAG_MTA1 construct (bases 1-400) showing the N-terminal 3X FLAG tag and positions of the PCR-validation primers (FLAG_FW and FLAG_Rev). Expected amplicon size is indicated. Bottom panel: PCR validation of FLAG_MTA1 transformants. A specific product is detected only in FLAG_MTA1, and absent in the wild-type parental strain (JRB510), the *mta1* mutant, and the no-template control (NTC). PCR was performed on total genomic DNA from asexually growing cells. NTC: no-template control; MW: molecular weight marker. **F**. Dot blot analysis of 10 ng RNase-treated genomic DNA from asexually growing *Oxytricha* strains of the indicated genotypes. Genomic 6mA is detected using an α-6mA antibody, and TEBP-β hybridization is used as a loading control. The wild-type strain is JRB310; *mta1* mutant lacks functional MTA1^7^; FLAG-MTA1 is a transgenic line expressing FLAG-MTA1 from the macronucleus in an *mta1* mutant background. **G**. Co-immunoprecipitation mass spectrometry analysis of FLAG-MTA1 using a FLAG antibody. Log2 values of normalized label-free quantification protein intensities of FLAG-MTA1 mated with JRB510 plotted against corresponding values from a wild type (strains JRB310 x JRB510) mating at 24h post-mixing. MTA1c components, which are all enriched in the FLAG-MTA1 sample, are highlighted in red. **H.** Immunofluorescence analysis of FLAG-MTA1 (magenta) subcellular localization 24h post-mixing of wild-type or FLAG-MTA1 x JRB510 strains, *n* = 7. DNA is labeled with DAPI (cyan). Z: zygotic nucleus (anlage).

**Figure S2. 6mA specifically marks retained DNA regions during development, related to Figure 2. A.** Meta-chromosome plots showing the proportion of 6mA [6mA/(6mA+dA)] in MDSs on 24h Precursor (top panel, for 1,272 chromosomes), 48h Partially-rearranged (middle panel, 21,702 chromosomes) or 60h Partially-rearranged (bottom panel, 19,702 chromosomes), relative to their MDS locations on the somatic chromosomes, plotting the first or last 1000 bp from telomere addition sites. The 6mA pattern in the parental macronucleus (0h post-mixing) is shown for reference (grey, 11,927 single-gene chromosomes). All chromosomes are aligned and oriented with their 5′ ends proximal to transcription start sites (TSSs). **B.** Abundance of 6mA called on individual SMRT-seq CCS reads, divided by total sequenced A/T base-pairs, for the indicated developmental stages and germline genome categories, as in Figure 2A. **C.** Proportion of genome coverage by SMRT-seq CCS reads (top panel) and of 6mA identified from individual CCS reads (bottom panel) overlapping with each of six functional germline genome categories at the indicated developmental stages. CCS coverage exclusively counts A/T base-pairs. “MIC” refers to the proportion of A/T across genome categories in the reference germline genome^23^. **D.** Enrichment of 6mA across six functional germline genome categories, calculated as the difference between the proportion of 6mA calls within each genome category (Figure S2C, bottom panel) and the proportion of total sequenced A/T bases in that category (Figure S2C, top panel), at the indicated developmental stages. Both 6mA counts and background A/T coverage were calculated from individual CCS reads.

**Figure S3. *mta1* mutant backcross has altered methylation during development, related to Figure 3. A.** Proportion of total SMRT-seq CCS reads assigned to each read bin (product, precursor, partially-rearranged, and unmapped reads) in *mta1* backcrosses at the indicated developmental timepoints in hours (see Table S1 for read counts). **B.** Meta-chromosome plots showing the proportion of 6mA [6mA/(6mA+dA)] at 0h in the parental somatic genome of wild-type (gray, 11,927 chromosomes) vs. *mta1* backcrosses (pink, 8,438 chromosomes). Plotted are the first or last 1000 bp from chromosome ends. Single-gene chromosomes are aligned and oriented with their 5′ ends proximal to transcription start sites (TSSs). **C.** Meta-chromosome plots showing the proportion of 6mA [6mA/(6mA+dA)] in the developing nucleus of wild-type (blue) vs. *mta1* backcrosses (pink) for 24h precursor (upper panel: blue,1,272 wild-type and pink, 1,265 *mta1* backcross single-gene chromosomes) and 60h partially-rearranged molecules (lower panel: blue, 19,702 wild-type and pink, 19,002 *mta1* backcross single-gene chromosomes) relative to their MDS locations on the somatic chromosomes. Plotted are the first or last 1000 bp from telomere addition sites. All chromosomes are aligned and oriented with their 5′ ends proximal to transcription start sites (TSSs). **D.** Enrichment of 6mA across six functional germline genome categories in wild-type vs. *mta1* backcrosses at the indicated developmental stages. Enrichment is calculated as log_2_[6mA(%) / A/T(%)], where %6mA is the proportion of 6mA within a genomic category and %A/T is the proportion of total sequenced A/T bases in that category. Both 6mA counts and background A/T coverage derive from pooled SMRT-seq subreads, excluding genomic sequence with fewer than 20 subreads. WT: wild-type. **E.** Proportion in the *mta1* backcross of genome coverage by SMRT-seq CCS reads (left panel) and of 6mA called on individual CCS reads (right panel) overlapping with each of six germline genome categories at the indicated developmental stages. CCS coverage exclusively counts A/T base-pairs. **F.** Enrichment of 6mA across six functional germline genome categories in the *mta1* backcross, calculated as the difference between the proportion of 6mA calls within each category and the proportion of total sequenced A/T bases in that category, at the indicated developmental stages. Both 6mA counts and background A/T coverage were calculated from individual CCS reads. **G.** MRM chromatograms of dmA as in Figure S1 in *mta1* backcrosses at indicated developmental timepoints; X-axis: retention time; Y-axis: mass response. **H.** Dot blot analysis of 25 ng RNase-treated genomic DNA from wild-type mating and *mta1* mutant backcrosses at the indicated hours post-mixing. Genomic 6mA was detected using an α-6mA antibody (top panel), and methylene blue staining was used as a loading control (bottom panel).

**Figure S4. Mutation of *mta1* leads to imprecise 6mA deposition in backcross, related to Figure 4. A.** Proportional MDS content of each CCS read plotted against the extent of 6mA methylation detected on that read, shown for 36h precursor reads from wild-type and *mta1* backcross SMRT-seq datasets. Values reflect CCS reads passing quality thresholds (≧ 20 subreads) and containing at least one 6mA site. Point color represents density of the data. The red line is a simple linear regression, and *r*^2^ is the square of the correlation coefficient. **B.** Strand bias of 6mA within individual SMRT-seq CCS reads at the 36h precursor developmental stage in wild-type (blue) and *mta1* backcrosses (red). Strand bias is calculated for each read as the absolute deviation from equal strand distribution: |(positive-strand 6mA) / (total 6mA) - 0.5|. Lower values indicate more symmetric strand deposition. **C.** Histograms of 6mA abundance across individual 36h CCS reads, shown for precursor reads (left panel) and partially-rearranged reads (“Partial,” right panel) in wild-type (blue) and *mta1* backcross (red) backgrounds. 6mA abundance is calculated per read as 6mA divided by all sequenced adenines in that read. The y-axis indicates the proportion of reads in each 6mA-abundance bin. **D.** Extent of methylation [6mA/(6mA +dA)], at single-molecule resolution, for 36h developmental stages at various subsets of well-covered 6mA sites. “Well-covered” includes only sites with CCS coverage ≥3 for the given developmental stage and genotype. Extremely-highly covered sites (CCS coverage >10) are also excluded. “All 6mA sites” includes all adenine positions methylated in at least one CCS read from any 36h read category (wild-type or *mta1* backcross). “In-category 6mA sites” includes sites that were found to be methylated in at least one CCS read within for the specific developmental stage and genotype read category. “Wild-type sites” include sites that were methylated in at least one CCS read from any wild-type 36h developmental read category. **E.** Snapshot of a representative region within a single MDS showing the methylation status of every ApT motif in individual partially-rearranged SMRT-seq CCS reads at 36h. Only reads spanning the full region and supported by ≥20 subreads are shown. Each dot represents the methylation state of an ApT motif on that read: fully methylated (black), hemimethylated on the sense strand (red) or antisense strand (blue), or unmethylated (gray).

**Figure S5. *mta1* mutant backcross exhibits a developmental delay, related to Figure 5. A.** The frequency of junctions surrounded by neither product- nor precursor-like sequence, referred to here as “other”, for the shared set of covered junctions at 36h (70,439) for partially-rearranged molecules in wild-type (blue) vs. *mta1* backcross backgrounds (pink). This feature is called “Other index” and defined as: “other” reads / (product-like reads + precursor-like reads + “other” reads). **B.** Proportion of RNA-seq reads mapping to the MAC or MIC reference genomes, or to neither, in all wild-type and *mta1* backcross replicates at 48h and 55h. (Read pairs were first mapped to the MAC genome, and read pairs with a properly-paired alignment were classified as MAC-derived. Remaining reads were then aligned against the MIC genome. Reads that did not map to either genome were categorized as unmapped.) **C.** Proportion of MIC-mapping RNA-seq reads (defined in B) that map to the germline genome categories of TBE transposons (black) and MIC-limited genes (grey) in wild-type and *mta1* backcrosses at 48h and 55h post-mixing, just before most cells die in the mutant backcross. Bars show the average across replicates. **D.** Average 6mA density across 79 length-normalized “ideal” TBE transposons at 60h; 6mA density calculated from single-molecule SMRT-seq data. These 79 TBEs are classified as “ideal” based on the presence of 3 ORFs, of appropriate length and orientation, plus overall transposon length (as in Figure 5E). Single-molecule 6mA calls from partially-rearranged reads were aggregated into 50 bp bins and averaged across TBEs. Wild-type (blue) and *mta1* backcross (red) profiles are shown. The three TBE ORFs are illustrated in the schematic model below.

**Figure S6. 6mA is a PIWI protein-dependent, epigenetic mark for DNA retention, related to Figure 6. A.** Confirmation of the absence of Otiwi1 expression in the new zygotic MAC of 24h mutant exconjugants from a mating between two *otiwi1* mutant lines by immunofluorescence using a human anti-Piwi1L antibody. Wild-type 24h exconjugant cells were used as a positive control. Z: zygotic nucleus (anlage), pMAC: parental MAC. The scale bar is 10 µm. **B.** Snapshot of the retained 62bp IES within the germline locus of the *mta1* mutant, showing 6mA calls identified on two partially-rearranged DNA molecules, identified via SMRT CCS reads that span the retained IES in the *mta1* mutant backcross and display 6mA within the IES; no WT reads contained 6mA within the IES.

## ACKNOWLEDGMENTS

We thank Marko Jovanovic, Elizabeth Valenzuela, Andy Hanneman, Edwin Escobar, and Nan Dai for mass spectrometry; Richard D. Morgan and Yvette Luyten for PacBio sequencing, Vishnu Kumary for early sample prep; Derek M. Clay for generating the original *otiwi1* mutants, Jiawen Zhou for help with raw SMRT-seq reads processing, Tasha José for the model figure; Leslie Beh, Mariusz Nowacki, and Clément Carré for helpful discussion; Jeeva Abraham, Ryan Tran, and Sheela George for laboratory support, and all Landweber lab members for discussion. This work was funded by NIH grant R35-GM122555 to L.F.L. This study used the Confocal and Specialized Microscopy Shared Resource of the Herbert Irving Comprehensive Cancer Center at Columbia University, funded in part through NIH/NCI Cancer Center Support Grant P30CA013696.

## AUTHOR CONTRIBUTIONS

M.T.A., Y.F., and L.F.L. conceived the project. M.T.A., Y.F. and D.J.V. performed computational and experimental analysis for most figures and tables. M.T.A., D.J.V, and L.F.L. wrote the manuscript, which Y.F., S.P., and W.E.J. edited. D.J.V. and Y.F. performed pooled subread SMRT-seq analysis. D.J.V. performed individual CCS-read SMRT-seq analysis. Y.F. designed and performed developmental index analysis. M.T.A., E.A, and P.E. performed microscopy. M.T.A. and P.E. performed DNA mass spectrometry. M.W.L. designed, generated, and characterized FLAG-MTA1 and performed MS and data analysis. E.A. genotyped FLAG-MTA1. M.T.A. and E.A. genotyped *otiwi1* mutants. E.A. validated *otiwi1* mutants and performed immuno-Southern hybridization using these lines. M.T.A. and M.W.L. performed RNA-seq, which M.T.A and D.J.V. analyzed. M.T.A, E.A., S.S. and P.E. performed SMRT-seq. Y.F. processed raw SMRT-seq data. S.P. and W.E.J. contributed critical discussion.

## RESOURCE AVAILABILITY

### Lead Contact

Further information and requests for resources should be directed to L.F.L.

### Materials Availability

No new unique reagents were generated in this study.

### Data and Code Availability

- PacBio SMRT-seq data are deposited in SRA under BioProject accession number PRJNA1372786 and NCBI GEO under accession number GSE313269. Illumina RNA-seq data are deposited in NCBI GEO under accession number GSE312406 and SRA under BioProject accession number PRJNA665991.
- Any additional information required to reanalyze the data reported in this paper is available from the lead contact upon request.

| REAGENT or RESOURCE | SOURCE | IDENTIFIER |
| --- | --- | --- |
| Monarch Genomic DNA Purification Kit | New England Biolabs | T3010 |
| NucleoSpin Tissue Genomic DNA extraction Kit | Macherey-Nagel | 740952 |
| Qubit dsDNA BR Assay Kit | Thermo Fisher Scientific | Q32853 |
| SYBR Gold Nucleic Acid Gel Stain (10,000×) | Invitrogen | S11494 |
| Ethidium Bromide Solution | Fisher Scientific | BP1302 |
| 10× TBE Buffer | Fisher Scientific | BP1333 |
| 10× TAE Buffer | Fisher Scientific | BP1335 |
| 20× SSPE Buffer | Fisher Scientific | BP1334 |
| Amersham Hybond N+ membrane | Cytiva | RPN303B |
| Amersham Hybond-XL membrane | Cytiva | RPN303S |
| Anti-N6-methyladenine antibody (rabbit) | Synaptic Systems | Cat# 202003; RRID: AB_2732849 |
| Goat Anti-Rabbit IgG (H+L)-HRP Conjugate | Bio-Rad | Cat# 170-6515; RRID: AB_2617112 |
| Tween-20 | Sigma-Aldrich | P9416 |

## METHODS

### Protein Extraction and Quantification

Cells were lysed using the NucleoSpin TriPrep kit (Macherey-Nagel) according to the manufacturer’s instructions. Total protein concentrations were determined using the Qubit Protein Assay (Thermo Fisher Scientific), and 10 µg of protein was loaded per lane for all western blots.

### SDS–PAGE and Total Protein Imaging

Protein samples were resolved on 4–20% Bio-Rad precast stain-free polyacrylamide gels. Following electrophoresis, total protein in each lane was visualized directly on the stain-free gel using a Bio-Rad imaging system to confirm equal loading.

### Membrane Preparation and Protein Transfer

Gels and blotting papers were equilibrated for 30 min in Bjerrum Schafer–Nielsen transfer buffer (48 mM Tris, 39 mM glycine, 20% methanol, 1.3 mM SDS). PVDF membranes (Bio-Rad, 1620177) were cut to size, activated in 100% methanol for 1 min, and equilibrated in transfer buffer. Semi-dry transfers were assembled on the blotting apparatus (two sheets blotting paper, PVDF membrane, gel, two sheets blotting paper, Trans-Blot SD Transfer Cell), removing air bubbles at each step. Transfers were performed at 15 V for 15 min at room temperature.

### Western blotting

Following transfer, membranes were washed three times for 15 min in TBST (20 mM Tris-HCl pH 7.6, 150 mM NaCl, 0.1% Tween-20) and blocked for 1 h at room temperature in 2% nonfat dry milk in TBST. Membranes were incubated overnight at 4°C with primary antibodies diluted 1:1,000–1:5,000 (Custom made Rabbit anti-TBE antibody^47^) in blocking buffer, washed three times in TBST, and incubated for 1 h with HRP-conjugated secondary antibodies diluted 1:5,000 (Goat Anti-Rabbit IgG (H + L)-HRP Conjugate #1706515). After an additional three 15-min washes in TBST, membranes were incubated for 5 min with ECL substrate (mixed 1:1 from components A and B, Clarity Western ECL Substrate, Bio-Rad) and imaged on an Amersham ImageQuant 800 chemiluminescence system. Precision Plus Protein™ Unstained Protein Standards (Bio-Rad, 1610363) and Precision Plus Protein™ Dual Color Standards (Bio-Rad, 1610374) were used as molecular weight markers.

## REAGENT SETUP

### SDS–PAGE Running Buffer (Tris–Glycine–SDS, 1×)

25 mM Tris, 192 mM glycine, 0.1% SDS.

To prepare 10× stock: 30.3 g Tris base, 144.1 g glycine, 10 g SDS per liter; dilute 1:10.

### Bjerrum Schafer–Nielsen Transfer Buffer (1 L)

48 mM Tris, 39 mM glycine, 20% methanol, 1.3 mM SDS.

### 10× TBS (500 mL)

200 mM Tris, 1.5 M NaCl, pH 7.6.

### 1× TBST (1 L)

20 mM Tris, 150 mM NaCl, 0.1% Tween-20.

### Blocking Buffer

5% nonfat dry milk in TBST.

### Cell culture

Cells were cultured in Pringsheim medium (0.11mM Na_2_HPO_4_, 0.08mM MgSO_4_, 0.85mM Ca(NO3)_2_, 0.35mM KCl, pH 7.0) and fed with *Chlamydomonas reinhardtii* and *Klebsiella pneumoniae* as previously described^74^. Matings were performed by starving the compatible mating types, mixing the mating types, and diluting to a concentration of 5000 cells per milliliter in Pringsheim medium and plating the cells in 10cm plastic Petri dishes. Matings were assessed several hours after mixing mating types by calculating the percentage of paired cells per total cells. The following strains crosses were used in this study: JRB310^25^ and JRB510^36^, for wild-type crosses; JRB510 and *mta1* mutants, generated in ref.^7^ (Contig12701.0 g10712)^25^, for *mta1* mutant backcrosses; FLAG_MTA1 transgenic line (generated in this study, by introducing FLAG_MTA1 in *mta1* mutant) backcrossed to JRB510, otiwi1 mutant clones 1 mated to clone 2 (both generated in this study) for otiwi1 mating experiments. For clone generation, ssRNA was microinjected into mating cells at 12 hours post-mixing according to previously published protocols^31^. Post-injected cells were allowed to recover in Volvic water for 2 days before picking single cells and plating them in Volvic to establish clonal lines. Clonal lines were expanded for 7-10 days, genotyped and cyst stocks were prepared by starvation.

### Dot blot

Genomic DNA was extracted from vegetative cultures of *Oxytricha* cells using a NucleoSpin Tissue kit (Macherey-Nagel) for FLAG_MTA1 dot blot (Figure S1E), and using NucleoSpin TriPrep kit (Macherey-Nagel) for developmental 6mA dot blot (Figures 1E and S3H). These samples were subsequently RNase treated by diluting 200 ng of DNA into 50 μL of 5 mM Tris with the addition of 1:20 RNase Cocktail (Ambion) and 1:50 RNase H (Thermo Fisher) and incubated at 37°C for 1 hour. RNase-treated DNA was purified using a MinElute PCR Purification Kit (QIAGEN), eluted in nuclease-free water (Ambion). DNA was then denatured by incubating at 95°C for 10 minutes, then placed on ice for 5 minutes. 10ng (FLAG_MTA1 6mA dot blot) or 25ng (developmental 6mA dot blot) denatured DNA was spotted onto an Amersham Hybond N+ (GE Healthcare Life Sciences) positively charged nylon membrane in 2.5 μL spots and dried for 5 minutes. The DNA was UV crosslinked to the membrane using a UVC-515 Ultraviolet Multilinker (Ultra-Lum) set to 70000 mJ/cm^2^. After crosslinking, the membrane was blocked in 5% NFDM in TBST for 1 hour at room temperature. The membrane was incubated with a 1:1000 dilution of α-m6A antibody (Synaptic Systems, 202003) in NFDM overnight at 4°C for FLAG_MTA1 activity in asexually growing cells (Figure S1E) or 1:3000 dilution for developmental 6mA assays, when 6mA is upregulated (Figures 1E and S3H). After primary incubation, the membrane was washed 3 times in TBST for 10 minutes, then incubated with a 1:3000 dilution of HRP-conjugated goat anti-rabbit antibody in NFDM for 1 hour. The membrane was washed 3 times in TBST for 10 minutes, then incubated with Clarity Western ECL Substrate (Bio-Rad) for 5 minutes, and imaged using an Amersham AI600RGB. DIG-labeled TEBP-ꞵ probes^75^ generated using a PCR DIG Probe Synthesis Kit (Roche) were used as loading control for the FLAG_MTA1 6mA dot blot, and methylene blue (MRC, MB 119) for 6mA dot blot in development.

### Genomic DNA extraction

Genomic DNA was extracted from *Oxytricha trifallax* mating cultures at 0, 12, 24, 36, 48, and 60h post-mixing using the Monarch Genomic DNA Purification Kit (New England Biolabs) for all SMRTseq experiments and DNA immunoblots, following the manufacturer’s instructions. Genomic DNA was extracted from *Oxytricha trifallax* mating cultures at 0, 12, 24, 36, 48, and 60h post-mixing using the NucleoSpin TriPrep kit (Macherey-Nagel) for developmental 6mA dotblots according to the manufacturer’s instructions. DNA was eluted in nuclease-free water and quantified using a Qubit fluorometer (Thermo Fisher Scientific). Genomic DNA for 6mA detection in *mta1* backcross lines and *otiwi1* mutants, as well as the corresponding wild-type cross JRB310 to JRB510, was prepared in the same way and quantified by Qubit prior to loading.

### Agarose gel electrophoresis

The 1 Kb Plus DNA ladder (#10787018, Thermo Fisher, Waltham, MA, USA) was used as a size standard. For developmental time-course immunosouthern analysis, 10 µg of undigested genomic DNA were separated on 1% agarose gels cast and run in 1× TBE (89 mM Tris-borate, 2 mM EDTA, pH ∼8.3). Gels were run until the dye front (New England Biolabs (NEB) Gel Loading Dye, Purple (6X) (B7024S) migrated near the bottom of the gel (∼300-500 bp). After electrophoresis, gels were stained with SYBR™ Gold Nucleic Acid Gel Stain (Invitrogen) and imaged under UV illumination to document equal loading and the integrity of high molecular weight DNA.

For developmental time-course immunosouthern analysis, 10 µg of undigested genomic DNA were separated on 1% agarose gels cast and run in 1X TBE (89 mM Tris-borate, 2 mM EDTA, pH ∼8.3). Gels were run until the DNA had migrated close to the bottom of the gel. After migration, gels were incubated with SYBR™ Gold Nucleic Acid Gel Stain (Invitrogen) and imaged under UV illumination to document equal loading and the integrity of high molecular weight DNA.

For immunosouthern analysis of *otiwi1* mutants and corresponding wild-type samples, 2 µg of genomic DNA collected at 0h post-mixing or at the 24h developmental timepoint were resolved on 0.8% agarose gels in 1X TAE (Fisher Scientific) supplemented with ethidium bromide (0.5 µg/ml). Gels were imaged under UV immediately after migration to verify loading and DNA quality.

### Gel treatment and DNA transfer to nylon membranes

Following electrophoresis, gels were incubated for 7 min in diluted HCl to 0.228M (4 mL of 36% HCl in 200 mL Milli-Q water) with gentle agitation to depurinate the DNA, then briefly rinsed in water. Denaturation was carried out through two sequential 30-min incubations in Southern denaturation buffer (1.5 M NaCl, 0.4 M NaOH), followed by a 15-min rinse in 20X SSPE (Thermo Fisher). DNA was transferred overnight by capillary blotting in 20X SSPE onto positively charged nylon membranes, using Amersham Hybond N+ (Cytiva) for developmental time-course samples and Amersham Hybond-XL (Cytiva) for *otiwi1* and corresponding wild-type samples. After transfer, membranes were air-dried on Whatman paper and genomic DNA was crosslinked by UV irradiation.

### Immunosouthern detection of 6mA

Membranes were equilibrated in TBST (20 mM Tris-HCl pH 7.6, 150 mM NaCl, 0.1% Tween-20) and blocked for 1h at room temperature in 5% nonfat dry milk in TBST. Detection of developmental 6mA dynamics and 6mA levels in *mta1* backcrosses and *otiwi1* mutants was performed using a rabbit anti–N⁶-methyladenine antibody (Synaptic Systems) diluted 1:3500 in blocking buffer and incubated overnight at 4°C with gentle agitation.

The following day, membranes were washed 3X 10-15 min in TBST and incubated for 1h at room temperature with an HRP-conjugated anti-rabbit secondary antibody (Bio-Rad) diluted 1:10,000 in blocking buffer. After three additional TBST washes, membranes were incubated for 5 min with chemiluminescent HRP substrate and imaged on a digital chemiluminescence detection system.

Signal intensities were used to assay the relative 6mA levels across developmental stages and between wild-type, crosses and *mta1* backcrosses, or *otiwi1* mutant crosses, respectively.

### Artificial nanochromosome construction and transformation

Genomic DNA from JRB310 was used as a template for PCR to amplify the portion of the MAC chromosome containing the coding and regulatory regions of MTA1 using primers 12701 5’ & 12701 3’. This amplicon was cloned using the TOPO XL-2 cloning kit (Invitrogen), and the resulting plasmid was modified to add a 3xFLAG tag to the N-terminus of the MTA1 coding region using the Q5 Site-Directed Mutagenesis kit (New England Biolabs) with primers MTA1 3xF SDM F & MTA1 3xF SDM R. This construct was PCR-amplified using primers 12701 5’ telo & 12701 3’ telo that add the double-stranded *Oxytricha* telomeric repeats to the 5’ and 3’ ends, then purified through ethanol precipitation and resuspended in nuclease-free water (Ambion). The resuspended DNA was filtered through an Ultrafree 0.22 μm PVDF centrifugal filter (Millipore). This construct was injected into the MAC of vegetatively growing MTA1 IES+ cells at a concentration >1 μg/μL^76^. After injection, cells were separated and grown clonally in individual wells of a 24-well plate and PCR-screened for the presence of the 3xFLAG tag using primers 12701 5’ & 12701 3xF SDM R.

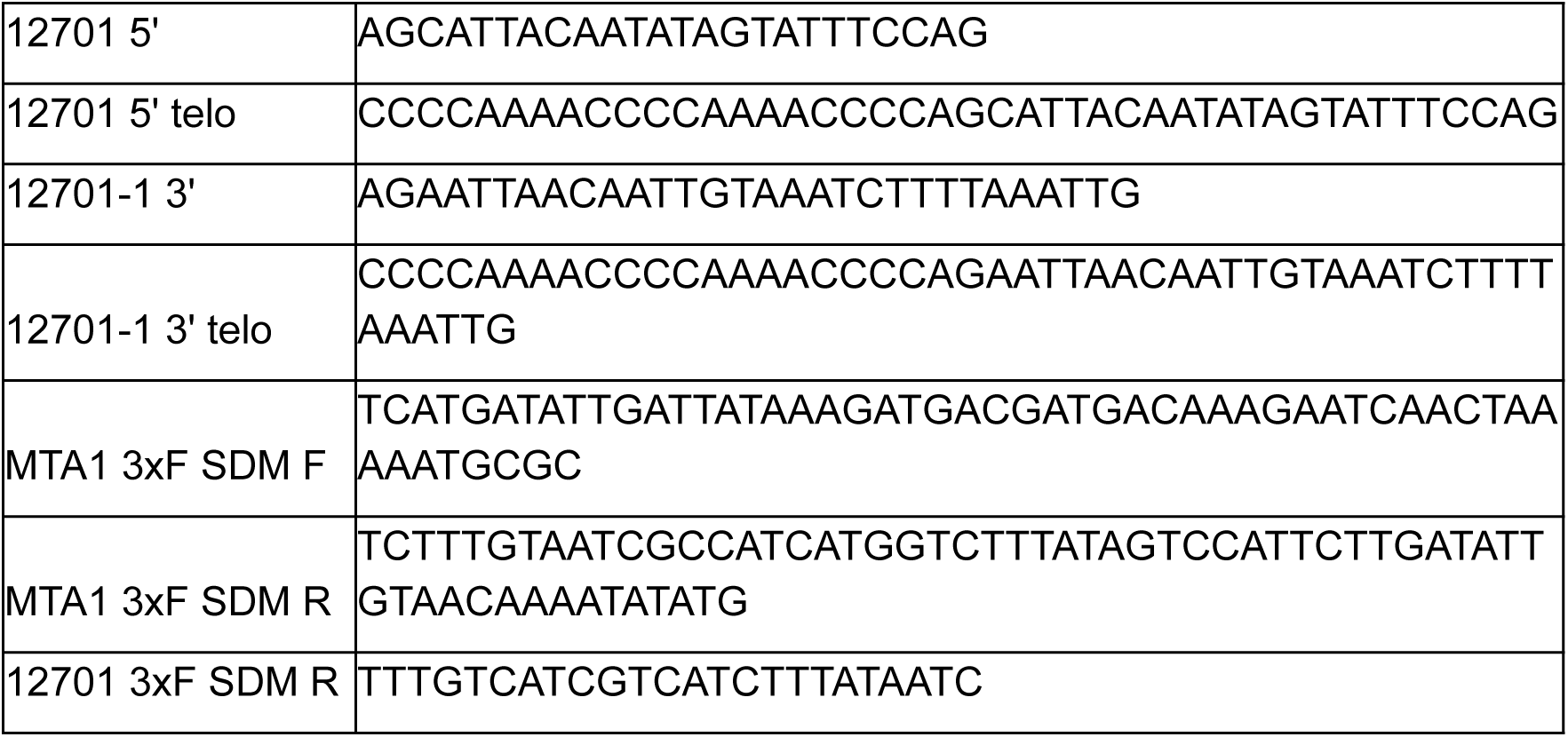

### Immunoprecipitation of FLAG-MTA1

Five million total cells of equal numbers of either JRB310 & JRB510 or FLAG-MTA1 & JRB510 were mixed after starvation and allowed to mate as described previously. The mating cultures were collected after 24 hours by concentrating on a nylon membrane and centrifugation at 130g for 1 minute. The cell pellets were snap frozen in liquid nitrogen, then thawed and suspended in Lysis buffer (20mM Tris-HCl pH 8, 135mM NaCl, 1mM MgCl2, 1mM EDTA, 1% NP-40, 10% glycerol) with the addition of Halt Protease and Phosphatase inhibitor (ThermoFisher). Lysates were cleared by centrifugation at 12000g for 15 min at 4°C. Cleared lysates were incubated with Dynabeads Protein G (Thermofisher) that had been bound to either 5 μg M2 anti-FLAG (Thermofisher) or mouse IgG for 3 hours at 4°C. Beads were washed twice with Wash buffer (50mM Tris pH 7.5, 150mM NaCl, 5% glycerol) + 0.05% NP-40, then twice with Wash buffer. After washing, the beads were incubated with 80μL urea/trypsin buffer (2M urea, 50 mM Tris pH 7.5, 1 mM DTT, 5 μg/mL MS grade bovine trypsin (Sigma-Aldrich)) at 25 °C for 1h shaken at 1000 rpm. The supernatant was removed and combined with two 60μL washes of the beads in urea/Tris (2M urea, 50mM TrisHCl pH 7.5). The proteins were reduced by incubation with 4mM DTT for 30 min at 25°C while shaking at 1000rpm, then alkylated with the addition of 10mM iodoacetamide and incubated for 45min at 25°C while shaking at 1000rpm. Samples were digested overnight by adding 0.5 μg Trypsin and incubation at 25°C while shaking at 700rpm. Digested peptides were acidified by addition of 1% formic acid and desalted using C18 StageTips.

### LC/MS-MS

LC/MS-MS on FLAG-MTA1 IP samples was performed using a Q Exactive HF coupled to a Waters nanoACQUITY UPLC. Peptides were separated at a flow rate of 400 nL/min using solvents A (0.1% formic acid in water) and solvent B (0.1% formic acid in acetonitrile) with the following gradients of solvent B: a 19 minute linear gradient from 2% to 10%, a 27 minute gradient from 10% to 22%, a 5 minute gradient from 22% to 30 %, a 4 minute gradient from 30% to 60%, and a 1 minute gradient from 60% to 90%. Samples were measured for 90 minutes in Data Dependent Acquisition mode and MS1 spectra were acquired at a resolution of 120000, an AGC target of 3e6, and scan range of 300 to 1800 m/z. The top 12 ions were chosen for MS2 analysis with a resolution of 15000, AGC target of 1e5, an isolation window of 1.6 m/z, and a normalized collision energy of 25.

100 ng of *Oxytricha* genomic DNA was digested using Nucleoside Digestion Mix (NEB # M0649S) according to manufacturer’s recommendations. LC-MS/MS analysis was performed in duplicate by injecting digested DNA on an Agilent 1290 UHPLC equipped with a G4212A diode array detector and a 6490A triple quadrupole mass detector operating in the positive electrospray ionization mode (+ESI). UHPLC was carried out on a Waters XSelect HSS T3 XP column (2.1 × 100 mm, 2.5 µm) with the gradient mobile phase consisting of methanol and 10 mM aqueous ammonium formate (pH 4.4). MS data acquisition was performed in the dynamic multiple reaction monitoring (DMRM) mode. Each nucleoside was identified in the extracted chromatogram associated with its specific MS/MS transition: dC [M+H]^+^at m/z 228→112, dG [M+H]^+^ at m/z 268→152, dT [M+Na]^+^ at m/z 265→149, dA [M+H]^+^ at m/z 252→136, and dmA [M+H]+ at m/z 266 →150. External calibration curves with known amounts of the nucleosides were used to calculate their ratios within the analyzed samples.

### Immunofluorescence

Images for developmental 6mA characterization were acquired using LSM 880 confocal microscope and Zen 2.1 software. 6mA and tubulin were detected using 458, 488, 514 nm multiline Argon laser and 561 nm DPSS laser, respectively. Nuclei were stained with DAPI and detected with 405 nm diode laser. Images for 6mA detection in *otiwi1* mutant background were acquired using Nikon Ti2 inverted microscope with an AXR resonant spectral scanning confocal microscope and NIS-Elements software at the [Confocal and Specialized Microscopy Shared Resource of the Herbert Irving Comprehensive Cancer Center at Columbia University]. Fluorophores were excited using a 405 nm diode laser (DAPI), a 488 nm diode laser (6mA), and a 561 nm DPSS laser (tubulin). For FLAG_MTA1 localization experiments and *otiwi1* mutants characterization fluorescent images were acquired on a Zeiss Axiovert 200 inverted microscope equipped with an Axiocam 506 monochrome camera. For immunofluorescence quantification, 6mA signal intensity was measured as the mean integrated pixel intensity per nucleus. No background subtraction or normalization was applied; raw pixel intensities were used for all analyses. For dual-channel composites, individual channels were pseudocolored and merged in Fiji^77^. 6mA intensity was plotted with Python matplotlib (Figure 1B and Figure 3B). Immunofluorescence was performed as previously described^31^. Primary antibodies used for immunofluorescence included anti-6mA (Synaptic Systems, Cat. No. 202003; 1:2000), anti-α-tubulin DM1A (Abcam, ab7291; 1:500), anti-PIWIL1/Otiwi1 (Abcam, ab12337; 1:350), and anti-FLAG (DYKDDDDK) Tag monoclonal antibody (Invitrogen™, clone FG4R, Cat. No. MA1-91878; 1:1000). Secondary antibodies included a 1:1000 Alexa Fluor™ 488–conjugated goat anti-rabbit secondary antibody (Invitrogen, A11034) and a 1:500 Alexa Fluor™ 546–conjugated goat anti-mouse secondary antibody (Invitrogen, A11030).

### PacBio library preparation

PacBio kit 2.0 was used to make DNA libraries according to manufacturer’s recommendations with minor modifications. First, 2 µg of *Oxytricha* genomic DNA (100 µl) were sheared using Covaris G tubes (Covaris # 520079) twice. We performed size fractions for each timepoint. For 0h, size fractions are 2-8kb and ≥9kb, For 24h and 36h, size fractions are 2-4kb, 4.5-8kb, 9-14kb and >14kb. For 48h and 60h, size fractions are 1-4kb and >4kb (Table S1, Table S2). 45 µl of NEBNext sample purification beads (NEB # E7103) were added and DNA was eluted in 48 µl with low TE buffer. 4 µl of DNA were run on 0.8% agarose gel to confirm that the size between 6 to 10 kb as recommended for PacBio library preparation and the rest was used for PacBio library construction. After DNA single strand overhang removal, DNA damage repair and end repair/dA tailing, SMRTbell ligation was performed for 4 hrs at 20°C and O/N at 4°C. After ligation, 2 µl of TLPK (NEB # P8111S) was added for 15 min at 37°C and inactivated for 10 min at 65°C. Nucleases mix (PacBio 2.0 kit) was added and incubated for 45 min at 37°C. A second 2 µl of TLPK was added for 15 min at 37°C to ensure nuclease degradation and inactivated for 10 min at 65°C. DNA clean-up was performed using 0.6x volume PacBio magnetic beads and DNA was eluted in 20 µl with low TE. After Qubit DNA quantification, DNA sample pool was made at 7.7 ng/µl. Primer was conditioned for 2 min at 80°C in 1x elution buffer before annealing to DNA pool for 30 min at 20°C. Polymerase loading was performed according to manufacturer’s recommendations and 120 µl of AMPure PB beads were added for 5 min at RT. The supernatant was discarded after placing the sample on a magnetic rack and beads were resuspended in adaptive loading buffer and incubated for 15 min at RT for DNA elution. The supernatant was kept on ice after placing the sample on a magnetic rack. Diluted internal control was added to 45 ng of pooled DNA library with adaptive loading buffer according to manufacturer’s recommendations. Libraries were loaded on PacBio Sequel II system.

### PacBio SMRT reads processing for 6mA annotations

We processed PacBio reads using command-line tools in SMRT^®^ Link v10.1. PacBio CCS reads for developmental timepoints (24-60h) were binned and assigned to four categories before downstream analyses: Product (MAC-matching), Precursor (MIC-matching), Partially rearranged (i.e., MIC-partially-matching) or Unmapped. For the analyzed developmental timepoints, PacBio CCS reads were first mapped to the reference *Oxytricha* MAC genomes of both strains: JRB310^25^ MAC and JRB510 MAC^36^ with pbmm2 (the SMRT^®^ Link wrapper for minimap2^78^). Perfectly MAC-mapped reads were considered as Product reads. The rest of the reads were then mapped to the reference *Oxytricha* MIC genome^27^. Perfectly MIC-mapped reads were labeled Precursor reads, while partially mapped reads were binned as Partially-rearranged reads (Figure S1B, Table S1). For 0h datasets, CCS reads were assigned as either JRB310 MAC, JRB510 MAC or JRB310 MIC based on their best BLASTN^79^ hit (Table S2). For each category, subreads were aligned to the MIC genome with pbmm2 and 6mA sites were annotated using ipdSummary (with parameters “--gff --identify m6A --methylFraction”). 6mA positions on the *Oxytricha* genome were visualized by IGV^80^. Final annotations used in this study only include 6mA sites which have subread coverage ≥20 and qv ≥20 (Table S3). For each developmental timepoint, CCS reads of wild-type mating and *mta1* backcross were subsampled so that they have equal read numbers in each category for 6mA calling (Table S1). 0h Product read dataset was not subsampled (Figure S3B). Note that the total number of reads are similar at 0h in wild type vs. *mta1*/+ backcross, but the *mta1*/+ backcross has more 510MAC-matching and unmapped reads, and wild-type has 29% more 310MAC-matching reads (Table S2, Figure S3B). For calculations of frequency of methylation (“6mA/(6mA+dA)”) from the pooled subread analysis, the underlying coverage of all A/T sites was calculated using aligned, pooled subreads for each read category. Excluding secondary alignments, bedtools (v2.29.2)^81^ genomecov was used to assess subread coverage of each base of the genome. Regions of the genome with minimum coverage of 20 subreads were extracted and converted to fasta files via bedtools getfasta, from which total coverage of A/T’s was counted. For calculations of frequency of methylation within germline genome sequence categories, bedtools intersect was used to subset the bed file containing regions with at least 20 subreads prior to counting A/T’s.

### Meta-chromosome analysis of 6mA sites on MDSs

To compare 6mA site distributions between developmental and vegetative growth stages, we converted the positions of 6mA sites on MDSs in the MIC genome to their corresponding positions in the MAC genome using a custom python script. After the position conversion, the MAC chromosomes were further filtered and processed to be included in plots: 1) they are one-gene complete MAC chromosomes with both telomeres; 2) they have at least one 6mA site called using the given read bin; 3) they are oriented by gene direction. Telomere sequences were clipped from the plots as in ref.^7^. The 6mA distribution of 24h partially-rearranged reads was found to match that of 0h Product reads; these 24h partially-rearranged reads were therefore excluded from further analysis on the basis of likely being derived from variants of nanochromosomes in parental macronuclei that did not perfectly map to the canonical version of the nanochromosome in the MAC assemblies.

### Single-molecule 6mA calling from individual PacBio SMRT reads

The presence of 6mA on each individual CCS read was measured by first splitting mapped subreads into separate .bam files on the basis of ZMW number using SMRT Link (v10.1) bamsieve, yielding individual .bam files for each CCS read. Only CCS reads with at least 20 subreads were retained for single-molecule 6mA calling. DNA modifications on each read were called by SMRT Link ipdSummary with parameters “--gff --identify m6A –methylFraction”. DNA modification calls were then filtered for exclusively 6mA with a minimum Qv threshold of 20. For each read, total counts and genomic locations of 6mA were recorded, and as were total counts and genomic locations of each A/T in the genome covered by the given read, excluding secondary alignments.

For each read, total 6mA and total genomic A/T coverage were recorded, both overall and within each of the six functional categories of the germline genome using bedtools (v2.29.2). 6mA calls were assigned to each functional category using bedtools intersect (with options -wa). Genomic A/T coverage within each functional category was measured using bedtools genomecov (with options “-bg -split -max 1”), followed by bedtools merge, to convert .bam files for individual CCS reads to .bed files, followed by bedtools intersect against genome category annotation files. 6mA strand bias of a read was measured by counting all 6mA calls on the positive and negative strands, then calculating the proportion of positive-strand 6mA out of total 6mA. The absolute value of the difference between this positive-strand proportion and 0.5 was defined as the “strand bias.” Penetrance of 6mA at a given genomic locus was measured by counting total CCS read coverage and total 6mA calls at that A/T site, then calculating the proportion of CCS reads bearing 6mA at that site. Average 6mA abundance on individual TBEs was calculated by measuring total coverage of A/T sites in that TBE using bedtools coverage, then summing total 6mA calls in that TBE. Individual TBEs were included in this analysis if cumulative coverage by CCS reads within the TBE amounted to at least 100 A/T base-pairs, and total depth of coverage of the TBE was less than 10x, to account for TBEs with extraordinarily high coverage, to which reads may have mapped ambiguously.

### Proportion of 6mA sites in MIC categories

Each nucleotide in the MIC genome is assigned to one of the categories with priority: MDS, TBE, MIC-limited genes, IES, Other repeats and Non-coding regions from ref. ^42^. For TBEs, we used subcategories from ref. ^46^.

### Rearrangement junction analysis

We investigated the genome rearrangement progress by estimating the status of rearrangement junctions using partially rearranged reads in both wild-type mating and *mta1* backcross. At the beginning of the rearrangement, the precursor junctions connect MDSs and MIC-limited regions. As the MIC-limited regions are removed from the genome, the product junctions only connect MDSs regions. A small proportion of junctions may not match the junctions captured in current MIC or MAC reference genomes, or could be errors in this biological process, and therefore would be categorized as “other”. For a clear definition of junctions, we focused on the 91,397 pointer junctions which have one-to-one correspondence between MIC and MAC genome. We first mapped each partially-rearranged CCS read to *Oxytricha*’s MAC and MIC genomes, and assigned the status of each junction as precursor-like, product-like or “other” by analyzing coverage using pysamstats (https://github.com/alimanfoo/pysamstats). We summarized the total number of reads in the three categories to characterize the status of each junction. We used two indices to characterize two perspectives in DNA rearrangement: 1) the Developmental index to characterize the progress. It is defined as product-like reads/(product-like reads+precursor-like reads), ranging from 0-1. 0 indicates the rearrangement has not started and 1 indicates the end. 2) the Other index to characterize possible errors. It is defined as other reads/(product-like reads+precursor-like reads+other reads). The plots only include pointer junctions that are covered by at least one read in both wild-type and *mta1* backcross datasets at the given timepoint.

### RNA-seq

RNA was extracted from frozen *Oxytricha* cell pellets using TRI Reagent (Ambion) according to manufacturer’s instructions. The RNA was DNase treated using a TURBO DNA-free kit (Invitrogen) using the rigorous DNase treatment protocol. For late developmental wild-type and *mta1* mutant backcross RNA-seq libraries, 1 µg of *Oxytricha* RNA was used for each timepoint (48 and 55h). Libraries were prepared using the polyA mRNA workflow of NEBNext® Ultra™ II Directional RNA Library Prep Kit for Illumina® (NEB # E7760S). Paired-end RNA-seq libraries were sequenced on an Illumina NovaSeq at New England BioLabs, with a read length of 2x75 bp. Adapters and low-quality sequences were trimmed with Trimmomatic (v0.39)^82^, with settings ILLUMINACLIP:TruSeq3-PE-2.fa:2:30:10, LEADING:3, TRAILING:3, SLIDINGWINDOW:4:15, MINLEN:36. Reads were aligned first to the 310 MAC reference genome with hisat2 (v2.2.1)^83^; read pairs that mapped concordantly to the MAC were labeled “MAC,” while unmapped read pairs and pairs that mapped only discordantly to the MAC were subsequently mapped to the 310 MIC reference genome. Discordant alignments to the MIC were allowed, given the presence of partially-rearranged DNA at these timepoints. FeatureCounts (v2.0.1)^84^ was used to call RNA-seq hits to the MAC and MIC transcriptomes (including TBEs, MIC genes, and transcribed germline-limited ORFs) with options “-p, -M, -O, --fraction, and -s 2” for the MAC transcriptome, and options “-p, -M, -O, and --fraction” for the MIC transcriptome. Given the predominance of reads deriving from previously-unidentified germline-limited transcripts, and the differences in MIC- vs. MAC mappability between the wild-type and *mta1* mutant backcross, counts-per-million-reads (CPM) were calculated for each previously-known germline transcript, and used as the basis for subsequent analyses. For comparisons of overall TBE and MIC gene transcript abundance, CPMs for all TBEs and CPMs for all MIC genes and transcribed germline-limited ORFs, respectively, were summed within each sample.

For analysis of RNA-seq reads derived from specific TBEs, the length and position of each ORF in each annotated TBE was assessed. Only TBEs with exactly one copy of each of the three ORFs, and in the canonical orientation, were retained. This set of TBEs was further filtered to include only TBEs with complete ORFs, where the length of each ORF was within 10% of the median length of that ORF within this subset. Maximum total TBE length was capped at 6kb, to remove TBEs with unusually long linker regions. The resulting set of 79 TBEs were termed “ideal,” and their length-normalized RNA-seq CPMs were plotted using deeptools (v3.5.1)^85^, with bin-size 50, normalized length 4000, and 100 base-pairs upstream and downstream included. Average density of 6mA calls across these same 79 complete TBEs were plotted in the same way.

Transcript abundance for MTA1, MTA9-B, and MTA9 described in Figure S1F was quantified from the polyA-selected RNA-seq dataset described in reference 41. TPMs were calculated using FeatureCounts (v2.0.1) from reads aligned *via* hisat2 (v2.2.1) to the MAC genes from the *Oxytricha trifallax* genome in ref. 27, with the following identifiers: MTA1 (Contig12701.0.0; g10712), MTA9-B (Contig17419.0; g11745), and MTA9 (Contig1237.1; g24932)). Transcript abundance was expressed as Transcripts Per Million (TPM). TPMs from biological replicates were averaged, with standard deviation reported. 0h, 12h, 24h, and 36h data points have three biological replicates, while the 48h timepoint has one biological replicate.

