## Supplemental Figures for "A PIWI protein-dependent DNA N6-adenine methylation pathway in *Oxytricha* protects genomic sequences from deletion"

**Figure S1: Adenine DNA methylation levels peak during development, related to Figure 1.**

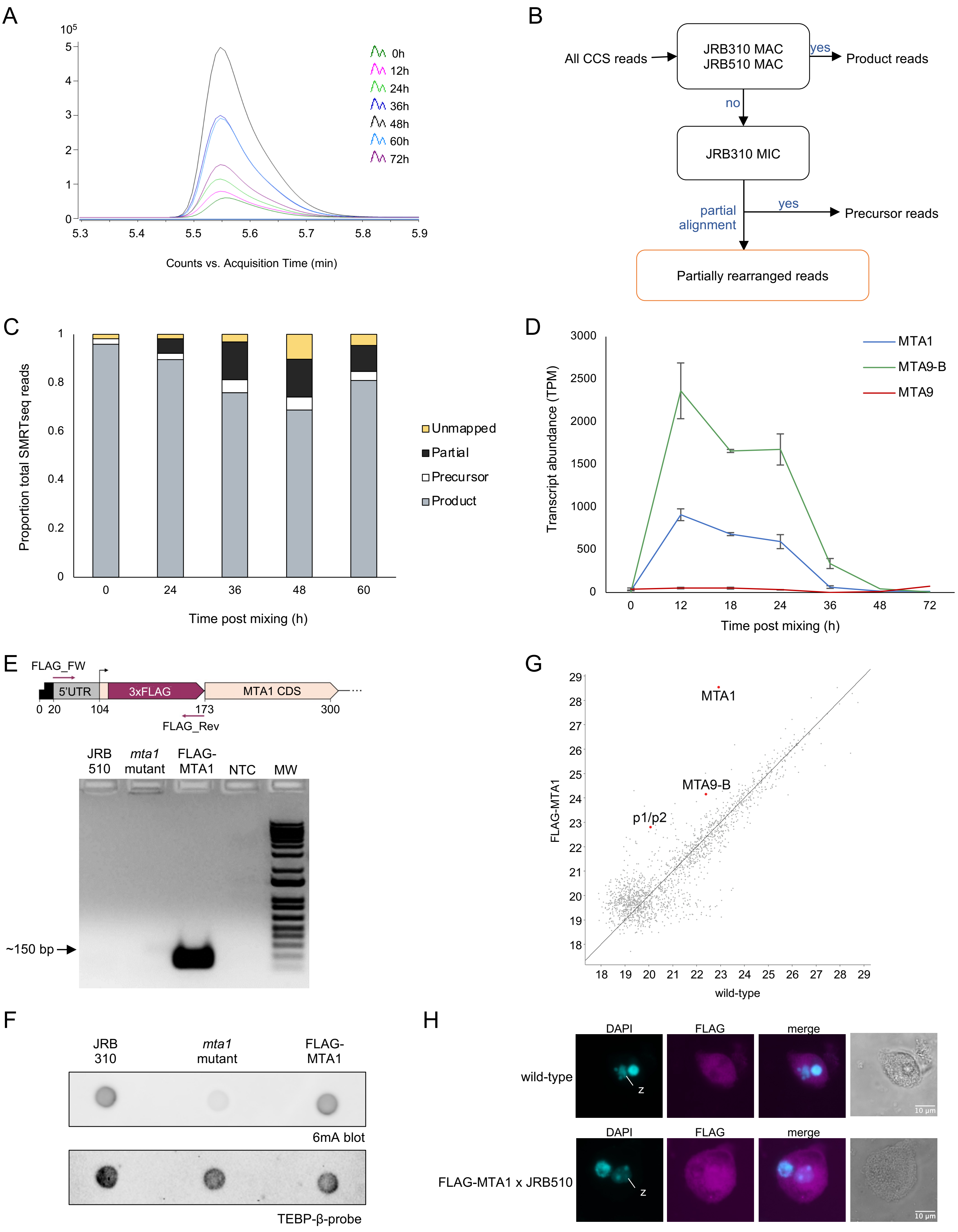

**Figure S2: 6mA specifically marks retained DNA regions during development, related to Figure 2.**

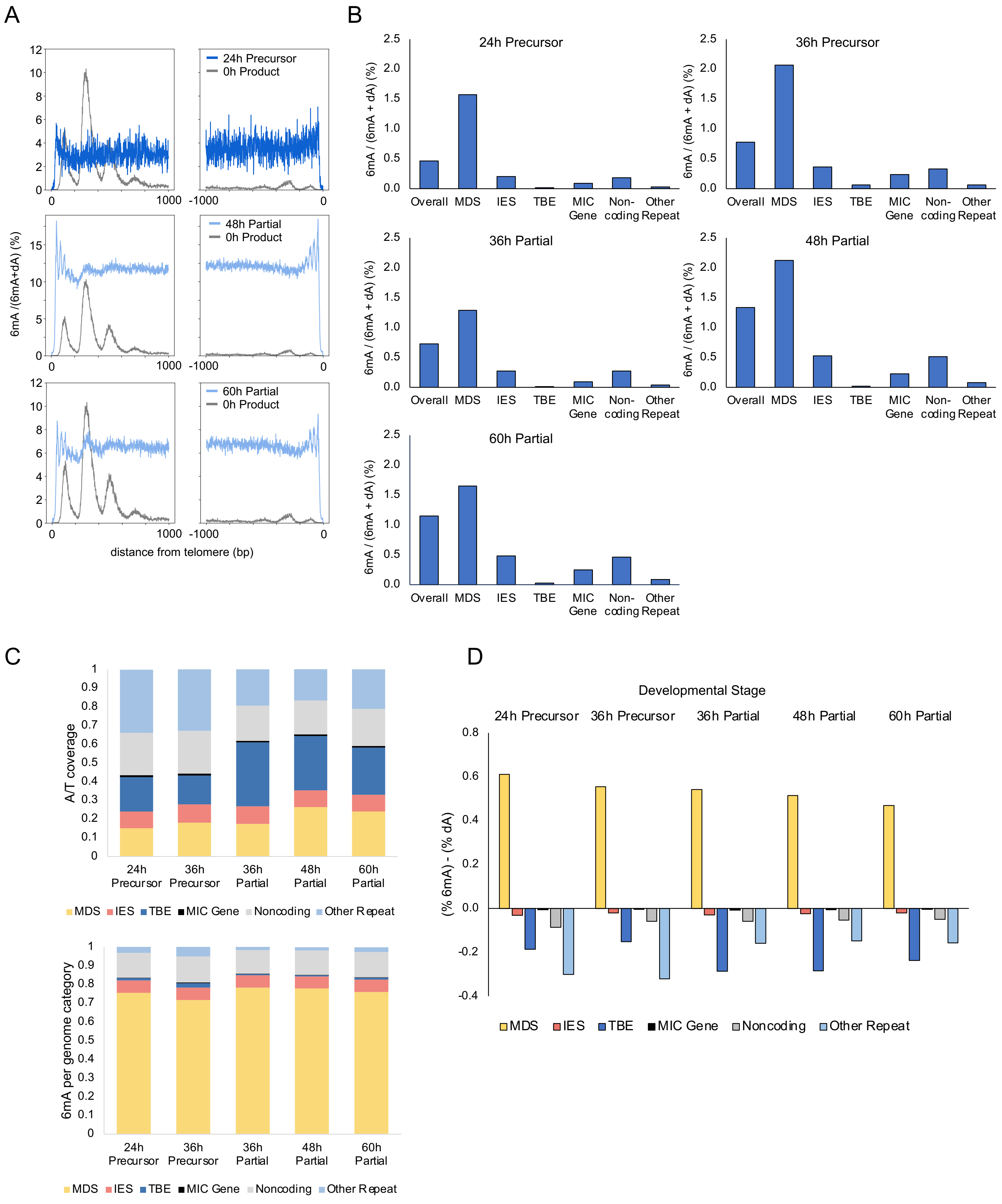

**Figure S3: *mta1* mutant backcross has altered methylation during development, related to Figure 3.**

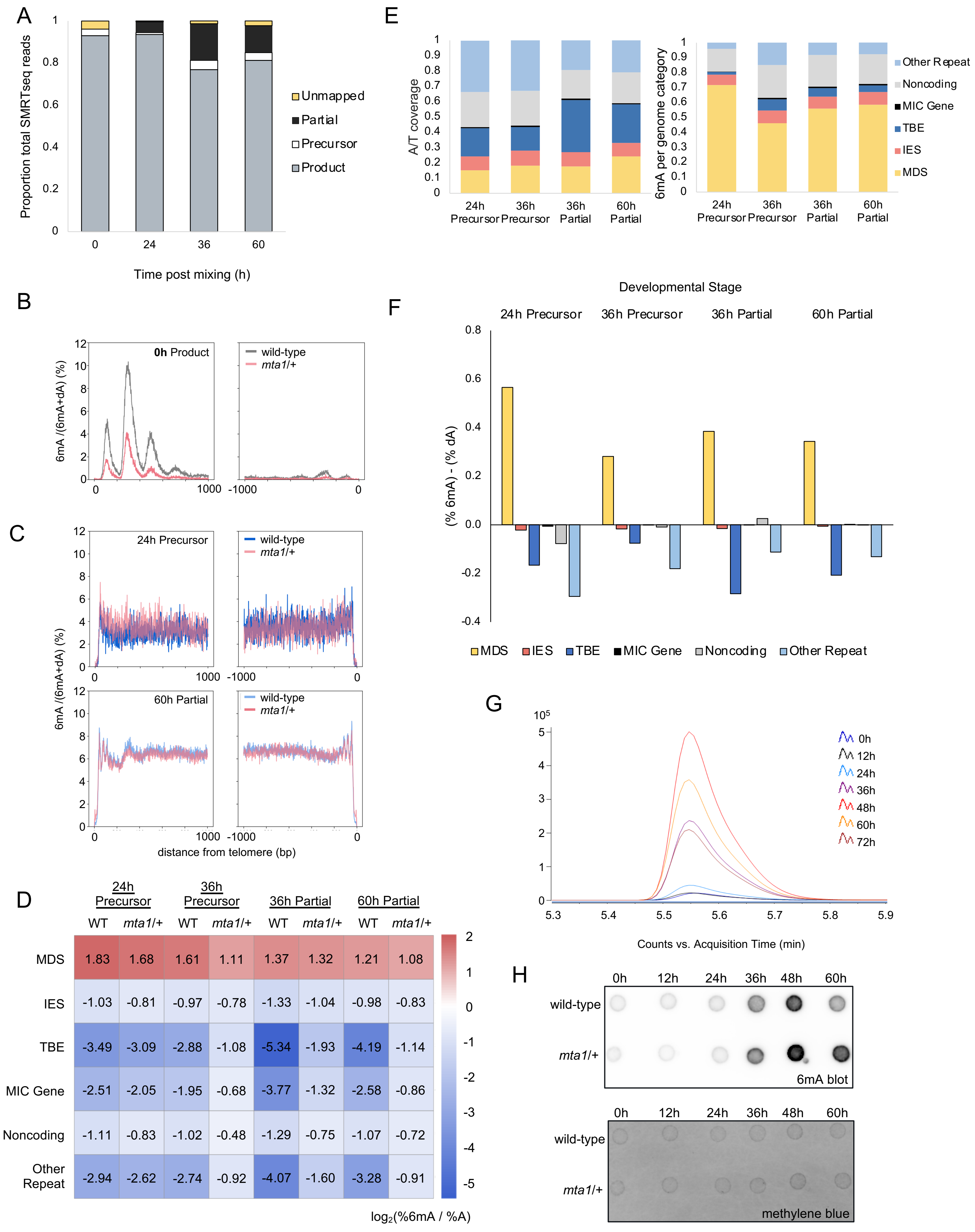

**Figure S4: Mutation of *mta1* leads to imprecise 6mA deposition in backcross, related to Figure 4.**

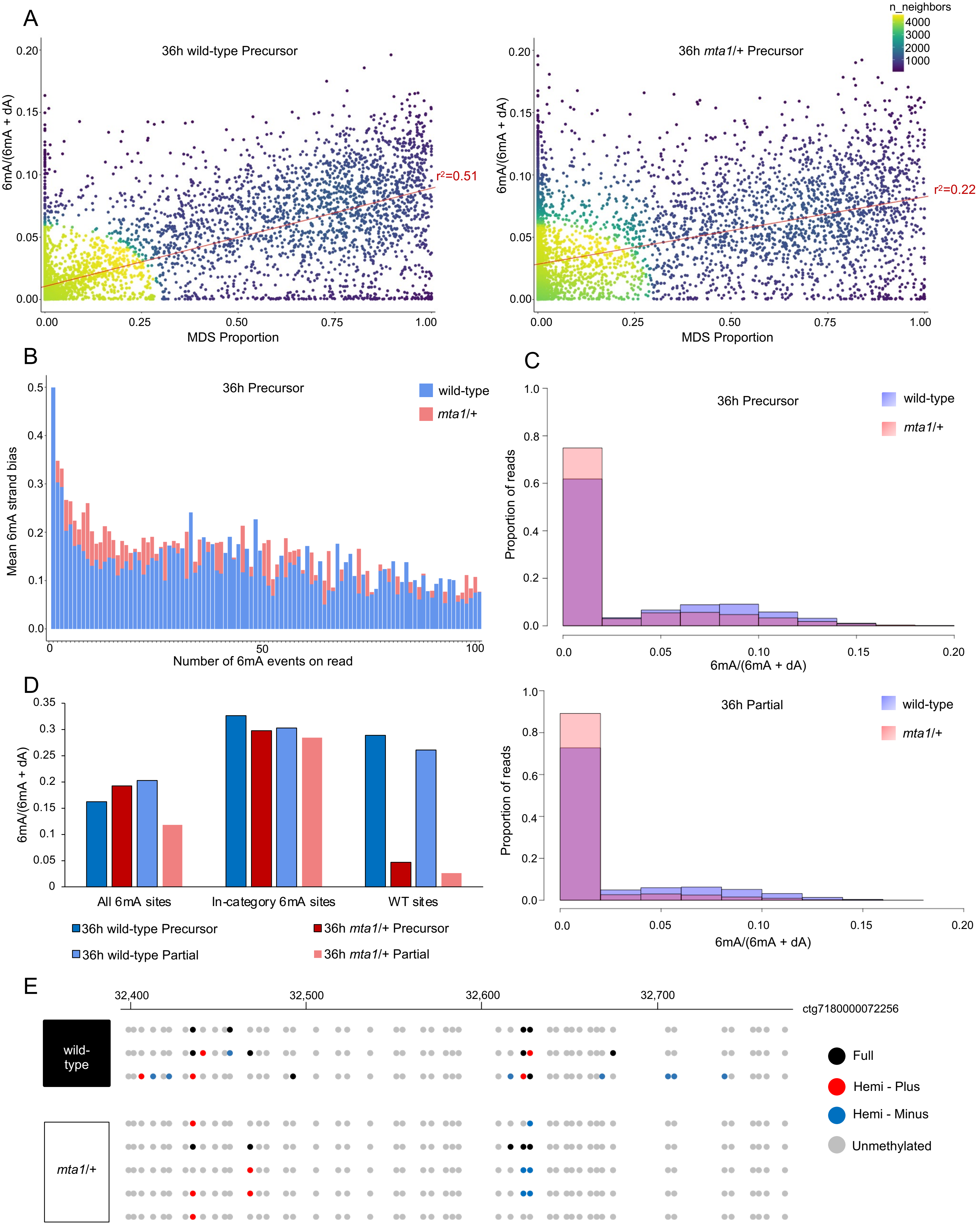

**Figure S5: *mta1* mutant backcross exhibits a developmental delay, related to Figure 5.**

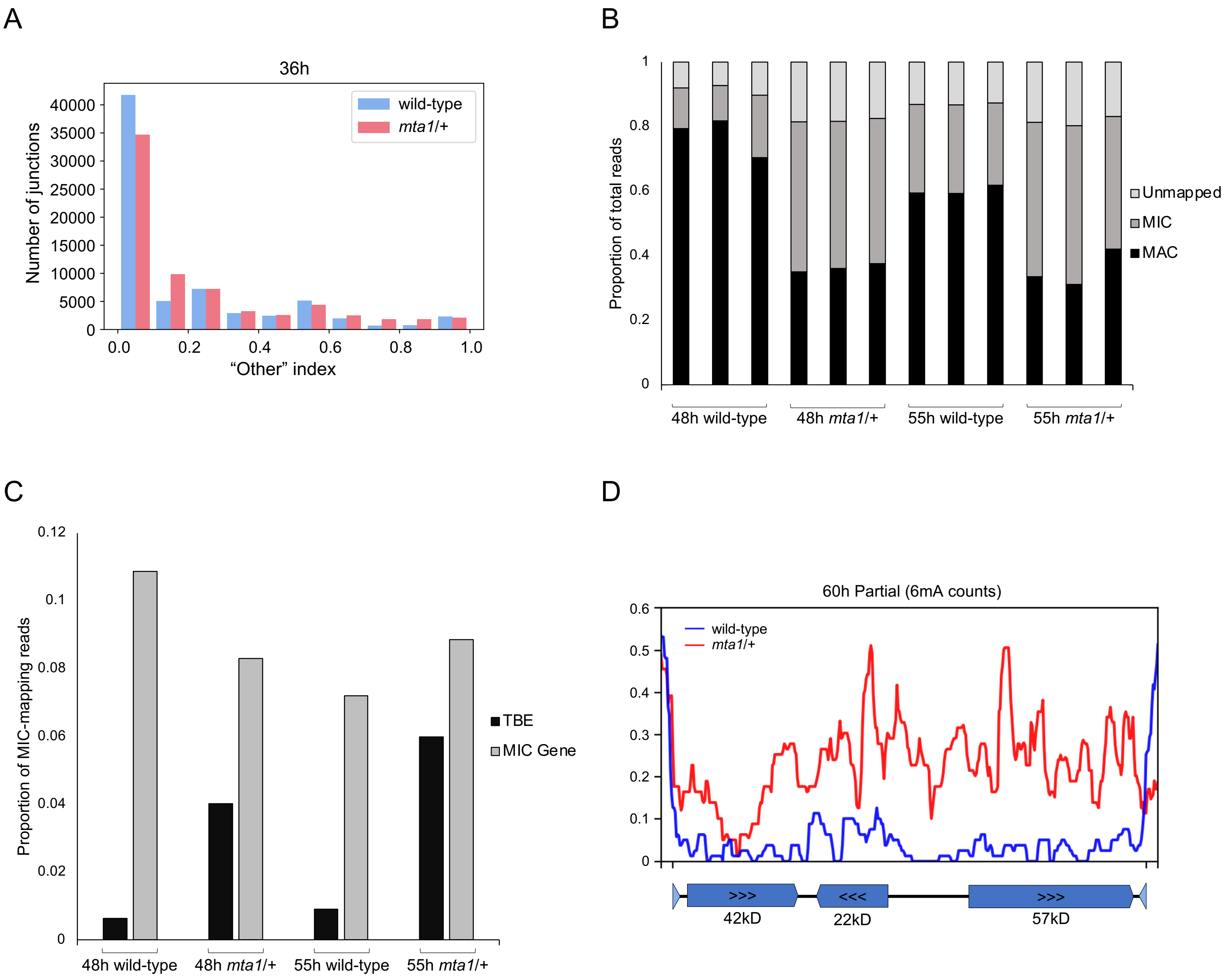

**Figure S6: 6mA is a PIWI protein-dependent, epigenetic mark for DNA retention, related to Figure 6.**

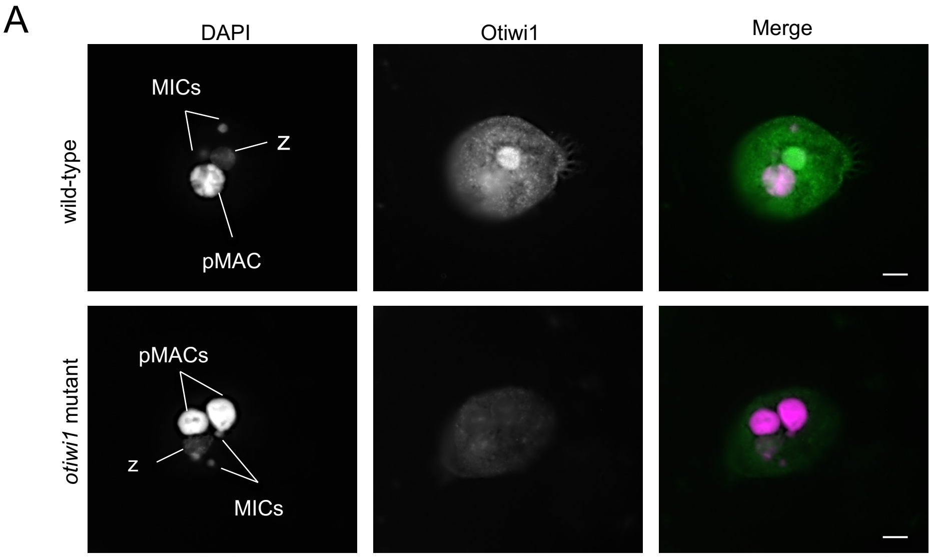


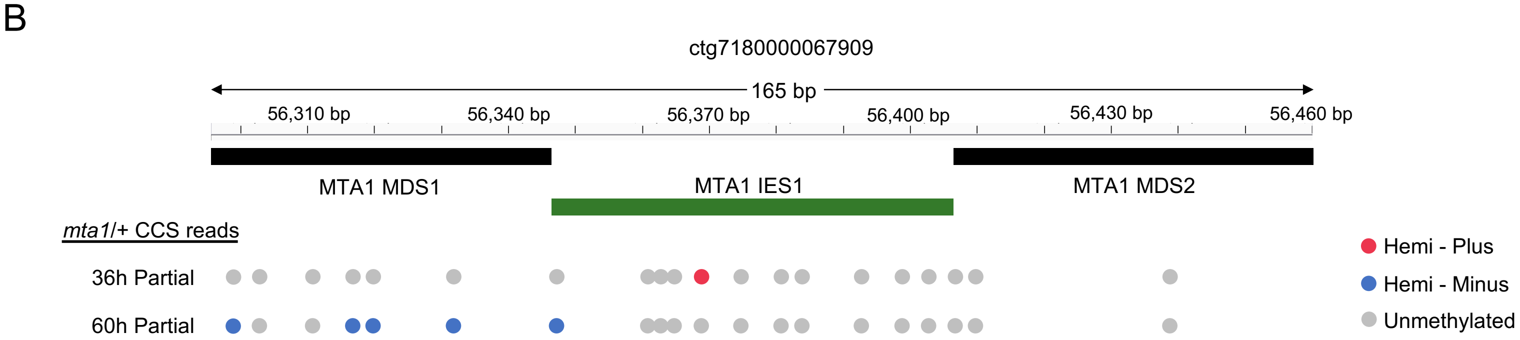
